# A Monte Carlo method for *in silico* modeling and visualization of Waddington’s epigenetic landscape with intermediate details

**DOI:** 10.1101/310771

**Authors:** Xiaomeng Zhang, Ket Hing Chong, Jie Zheng

**Author notes:** These authors contributed equally to this work (Xiaomeng Zhang, Ket Hing Chong). Correspondence: Jie Zheng.

## Abstract

Waddington’s epigenetic landscape is a classic metaphor for describing the cellular dynamics during the development modulated by gene regulation. Quantifying Waddington’s epigenetic landscape by mathematical modeling would be useful for understanding the mechanisms of cell fate determination. A few computational methods have been proposed for quantitative modeling of landscape; however, to model and visualize the landscape of a high dimensional gene regulatory system with realistic details is still challenging. Here, we propose a Monte Carlo method for modeling the Waddington’s epigenetic landscape of a gene regulatory network (GRN). The method estimates the probability distribution of cellular states by collecting a large number of time-course simulations with random initial conditions. By projecting all the trajectories into a 2-dimensional plane of dimensions *i* and *j*, we can approximately calculate the quasi-potential *U* (*x*^*i*^, *x*^*j*^) = −ln *P* (*x*^*i*^, *x*^*j*^), where *P* (*x*^*i*^, *x*^*j*^) is the estimated probability of an equilibrium steady state or a non-equilibrium state. A state with locally maximal probability corresponds to a locally minimal potential and such a state is called an attractor. Compared to the state-of-the-art methods, our Monte Carlo method can quantify the global potential landscape (or emergence behavior) of GRN for a high dimensional system. The same topography of landscape can be produced from deterministic or stochastic time-course simulations. The potential landscapes show that not only attractors represent stability, but the paths between attractors are also part of the stability or robustness of biological systems. We demonstrate the novelty and reliability of our method by plotting the potential landscapes of a few published models of GRN. Besides GRN-driven landscapes of cellular dynamics, the algorithm proposed can also be applied to studies of global dynamics (or emergence behavior) of other dynamical systems.

## Introduction

The Waddington’s epigenetic landscape has been recognized as a powerful metaphor for explaining the phenomena of embryonic development and cellular differentiation in biology^1,2,3,4^. The essence of the conceptual model proposed by Waddington is the ability to explain the emergent properties of cell fate decisions^5^. At least two types of approaches based on dynamical systems theory for quantifying the Waddington’s epigenetic landscape have been used. The first is the discrete formalism of Boolean network modeling^6,7^ and the second is continuous modeling in the form of ordinary differential equations (ODEs)^8,9,10^. This paper is focused on the second approach in the form of ODEs.

Recently, a few methods for quantifying and plotting Waddington’s epigenetic landscape based on gene regulatory networks (GRNs) have been proposed^11,12,13,14,15^. A key step in these methods is the formulation of a potential (or quasi-potential) value for the dynamical system of GRN that can be displayed as a landscape. For example, Bhattacharya *et al.*^11^ proposed a method for mapping aligned trajectories of the dynamical system of GRN in ODEs to a “quasi-potential” surface in the *x* -*y* phase space. When investigating mathematical models of two important processes in development, cell-fate induction and lateral inhibition^12^, Ferrell showed that the unique formulation of the potential surface for the lateral inhibition model can produce a pitchfork bifurcation, which is consistent with Waddington’s epigenetic landscape where a ball representing a cell is moving down the hill and then bifurcate into two valleys (i.e. two stable states).

Later, Zhou *et al.*^15^ proposed a theoretical framework for the decomposition of vector fields which enables the computation of a “quasi-potential function” for multi-attractor systems. Among the recent methods for quantifying Waddington’s epigenetic landscape is the one proposed by Li and Wang^13^ who used statistical mechanics to quantify the potential landscape through self-consistent mean field approximation. Their formulation of the potential based on the probability distribution of steady states captures the global potential landscape and the global barrier height measured by the “potential difference between the two attractor minimums and the saddle point on landscape”^13^. Li and Wang^13^ also used a path integral method to obtain the kinetic paths of transition between attractors. The self-consistent mean field approximation method has been implemented into a software package named “NetLand” by our group to facilitate the drawing of Waddington’s epigenetic landscape^16^.

Although the method proposed by Li and Wang can quantify potential landscapes for high dimensional GRNs, their method has limitation in the lack of realistic details of the landscape. Moreover, the high dimensionality of the GRN as measured by the number of genes poses challenges for modeling, analysis and visualization; for example, the methods proposed by Bhattacharya *et al.*^11^ and Ferrell^12^ allow two variables only. In this paper we propose a simple and novel Monte Carlo method for quantifying the Waddington’s epigenetic landscape of GRNs of more than two genes. Our algorithm projects the time-course trajectories into a 2-dimensional plane of dimensions *i* and *j* to calculate the probability distribution and potential *U* (*x*^*i*^, *x*^*j*^) = −ln *P* (*x*^*i*^, *x*^*j*^), where *P* (*x*^*i*^, *x*^*j*^) is the estimated probability of an equilibrium steady state or a non-equilibrium state. We demonstrate unique features of the proposed method by plotting the landscapes of a few case studies of GRN from two-dimensional to higher dimensional models. In our Monte Carlo method a large number of random initial conditions drawn from the state space are used to calculate time-series trajectories based on ODEs. A novelty of our method is the projection of the time-series trajectories into a plane that is divided into grid boxes to estimate the probability distribution and the quasi-potentials of cell states. The landscape altitude proportional to the quasi-potential, when laid out on the *x* -*y* plane, can capture detailed features of the dynamical system, such as basin of attraction, unstable manifolds connecting two attractors, spiral attractors and limit cycle attractors in the potential landscape.

Testing on a few published models of GRN showed that our Monte Carlo method can successfully quantify global potential landscapes consistent with the state-of-the-art methods. The case studies demonstrate the power of our computational method in uncovering the detailed dynamical behaviors of GRNs that other methods fail to capture. In addition, we have also used the stochastic approach of Chemical Langevin Equation (CLE) to estimate the probability distribution of states by collecting simulated time series. The potential landscape constructed by the stochastic approach turned out to be consistent with the landscape by the deterministic approach. Our analysis indicates that the structure of a GRN (or in other words model interactions) can contribute to the robustness of the attractors to noise. Moreover, we argue that in the Waddington’s epigenetic landscape not only the attractors represent stability, but that the paths between attractors characterised by unstable manifolds of saddle point or stable manifolds to attractor also contribute to the stability or robustness of gene regulatory systems for cell fate decision.

## Results

### Quantifying the potential landscape of non-equilibrium and equilibrium states

To demonstrate the capability of the proposed method, we selected four real models of GRN and one artificial network model. These models are: (1) a bistable synthetic toggle switch^17^, (2) a model of cancer attractors^18^, (3) an artificial network with spiral attractors^15^, (4) a cell cycle oscillator model^19^, and (5) a stem cell differentiation and reprogramming model^13^. For the model equations, readers may refer to the original papers or the source code included in our MATLAB package (see the Additional material). These examples illustrate that our method can capture distinct details of dynamical systems, e.g. attractor, repeller, unstable manifold, saddle point, spiral attractor and limit cycle attractor.

#### Example 1: Bistable synthetic toggle switch from Gardner *et al.*^17^

In the first example, our method is used to analyse the transient properties of a synthetic genetic toggle switch proposed by Gardner *et al.*^17^. The model equations and parameter values we used are the same as given by Segel and Edelstein-Keshet^20^. Phase plane analysis shows that the system displays two attractors and one saddle point, and the saddle point is formed by both stable and unstable manifolds^17,20,21^. According to the dynamical systems theory, stable manifolds are characterized by eigenvectors with negative real eigenvalues, whereas unstable manifolds are characterized by eigenvectors with positive real eigenvalues.

By applying the Monte Carlo method which is presented as Algorithm 1 in this paper (see Methods), we obtained the landscape as shown in Figures 1a-b. The potential landscape shows two attractors and there is a valley connecting the two attractors. By comparing the potential landscape with the phase plane, the valley (or kinetic path) in Figure 1a is found to be formed by the unstable manifolds of saddle point or stable manifolds of attractors, which cannot be generated by analysing steady states only. Between the two attractors, there is a saddle point (Figure 1b). It can be observed from the plotting of the 3D landscape that the unstable manifolds of saddle point form the valley, whereas the stable manifolds of the saddle point form the separatrix or boundary between the two attractors.

**Figure 1:**
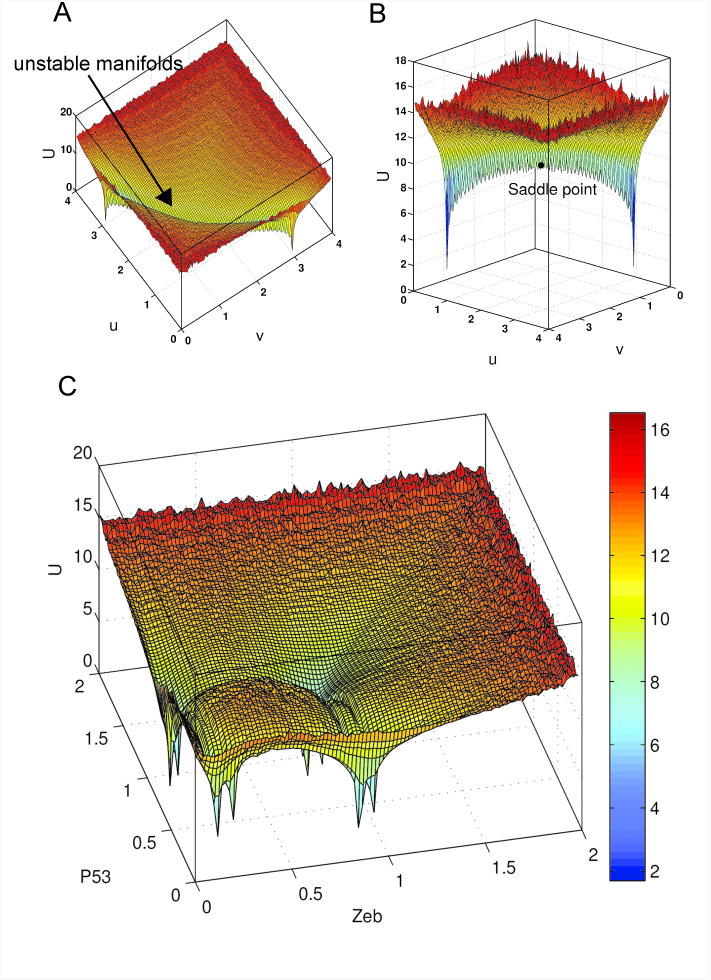
Potential landscapes for Example 1 and Example 2. **(a)** 3D view of a genetic toggle switch based landscape: The landscape displays two attractors that are connected with a kinetic path formed by unstable manifolds. **(b)** Side view of the genetic toggle switch based landscape in Example 1: The landscape displays two basins of attraction that are connected by the unstable manifolds. The saddle point separates the two attractors. The saddle point is also a tipping point (or barrier height) for transition along the kinetic path between the basins of attraction. **(c)** Validation of our method by reproducing the potential landscape of Li and Wang^18^ in Example 2. By comparing the location of the attractors labeled by Li and Wang^18^, we identified the attractors and their corresponding cell states. The landscape displays four main basins of attraction for normal cells, cancer cells, stem cells and cancer stem cells. However, our landscape is slightly different from the landscape plotted in the original paper (Li and Wang^18^): each of the main basins of attraction is divided into two attractors which are connected by kinetic paths. These kinetic paths may correspond to the kinetic paths suggested by Li and Wang^18^ where their results suggested that cell changes state according to different directions of the kinetic paths. The color corresponds to the value of the quasi-potential *U*.

This simple model of GRN shows that our method of formulating the potential can capture some transient properties of a dynamical system. For example, when there is a valley between two attractors in the potential land-scape, the kinetic path is formed by unstable manifolds. The saddle point sets a threshold for the barrier height that can separate the two attractors. The potential landscape displays a three-dimensional view of the phase plane and shows the attractors and saddle point more clearly than a two-dimensional plane only. In particular, the saddle point is shown to have one convex up (local minimum) and one concave down (local maximum) in the opposite directions. The result from this example suggests that not only attractors represent stability but the kinetic paths are also part of the stability of GRNs.

#### Example 2: Cancer attractors from Li and Wang^18^

To test if our method can handle a network of more than two genes, we choose a six-gene network model proposed by Li and Wang^18^. This gene network was reported to produce four attractors representing cancer stem cells, stem cells, cancer cells and normal cells^18^. The potential landscape of this model generated by our algorithm, however, shows four pairs of attractors in which the two attractors in each pair are close to each other (Figure 1c). Our timecourse data analysis of the trajectories end points (when simulation time end protein levels reached nearly unchanged values) confirmed that there are 8 attractors (data not shown). The result suggests that Li and Wang^18^ may have joined the two nearby attractors into one attractor and thus resulted in only four attractors.

This result indicates that the potential landscape obtained using our method can capture more details of attractors. In addition, the landscape also contains 5 valleys (Figure 1c). These valleys are formed by unstable manifolds, as discussed in the first example, and contain the information about saddle points as barrier heights for separating two attractors. These valleys are similar to the kinetic paths in Example 1 in illustrating the transition from one attractor to another attractor in the landscape.

#### Example 3: Spiral attractors from Zhou *et al.*^15^

The third example was selected to demonstrate the capability of our method to capture the dynamical features of spiral attractors. This model is an artificial network of two variables proposed by Zhou *et al.*^15^. The original landscape shows four attractors, four saddle points and one repeller in the center. The potential landscape generated using our method is shown in Figures 2a-b. There are four attractors on the four corners of the landscape and one repeller in the middle. However, the basins of attractor are unique in that they are formed by a counterclockwise spiral (Figure 2b). The landscape also shows four valleys (or kinetic paths) connecting the four attractors. The valleys are formed by the unstable manifolds of saddle points as discussed earlier.

**Figure 2:**
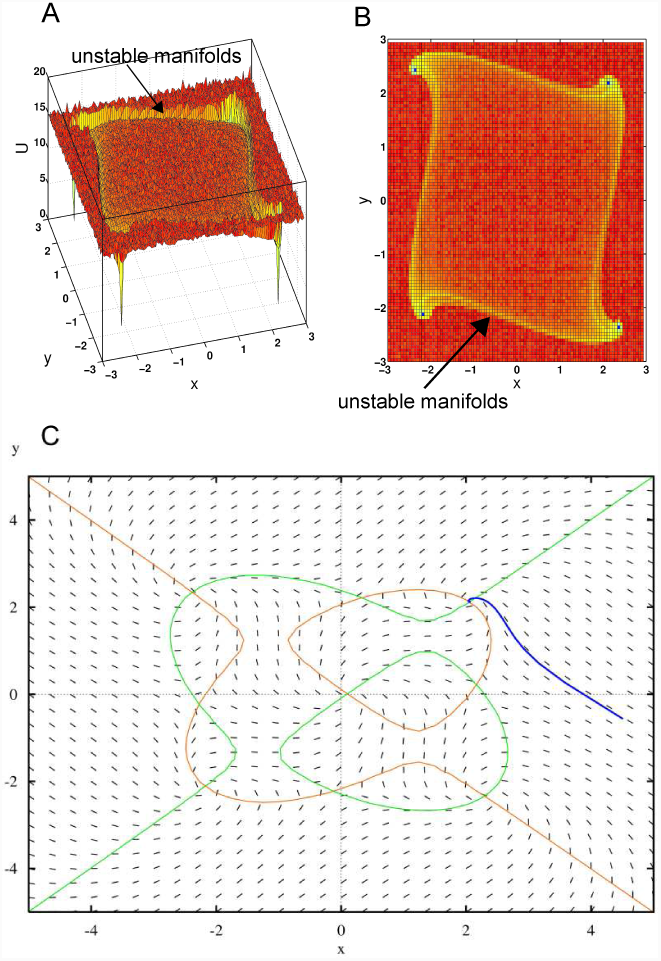
Potential landscape for Example 3. The Waddington’s epigenetic landscape displays four basins of attraction. **(a)** 3D view: The Waddington’s epigenetic landscape contains four spiral attractors (four blue dots) and the directions of the spirals are all counterclockwise. **(b)** Top view: There are unstable manifolds connecting any two attractors. **(c)** A conventional phase plane analysis shows the two nullclines where x or y does not change (green for y-nullcline and brown for x-nullcline) and the vector field. Intersections of the nullclines indicate the steady states. In this phase plane there are 9 intersection points corresponding to the 9 steady states: 5 unstable and 4 stable. The stability of the steady state can be determined by checking the eigenvalues of the eigenvectors: negative eigenvalues indicate stable steady states whereas positive eigenvalues indicate unstable steady states. Also shown is an example of a trajectory (blue line) being attracted to the stable steady state (one of the intersections between the nullclines at the top right). (Figure 2c was generated using XPPAUT, which can be downloaded from http://www.math.pitt.edu/~bard/xpp/xpp.html)

The conventional phase plane analysis can illustrate the vector field and the flows of the trajectories (Figure 2c), but it can only show the flows in terms of the counterclockwise spiral in a two-dimensional plane. Here, our potential landscape can quantify and depict the spiral attractors in a three-dimensional view of the potential landscape. The formulation of the potential *U* = −ln *P* which includes the probability of non-equilibrium state enables the quantitative description of the detailed transient behavior of the dynamical system.

#### Example 4: Cell cycle modeling by limit cycle oscillator from Ferrell *et al.*^19^

The fourth example is used to test the capability of our method to investigate another type of attractors, namely limit cycle attractor. This example is a 3-gene network model of limit cycle oscillator for modeling cell cycle control proposed by Ferrell *et al.*^19^. A few key proteins for controlling cell cycle were observed to oscillate^22^, and Ferrell *et al.*^19^ proposed an ODE model of gene network to explain why these proteins can oscillate in a limit cycle. Based on this 3-gene network we used our method to generate a potential landscape. The resulting potential landscape in Figures 3a-b shows a limit cycle attractor (in blue color).

**Figure 3:**
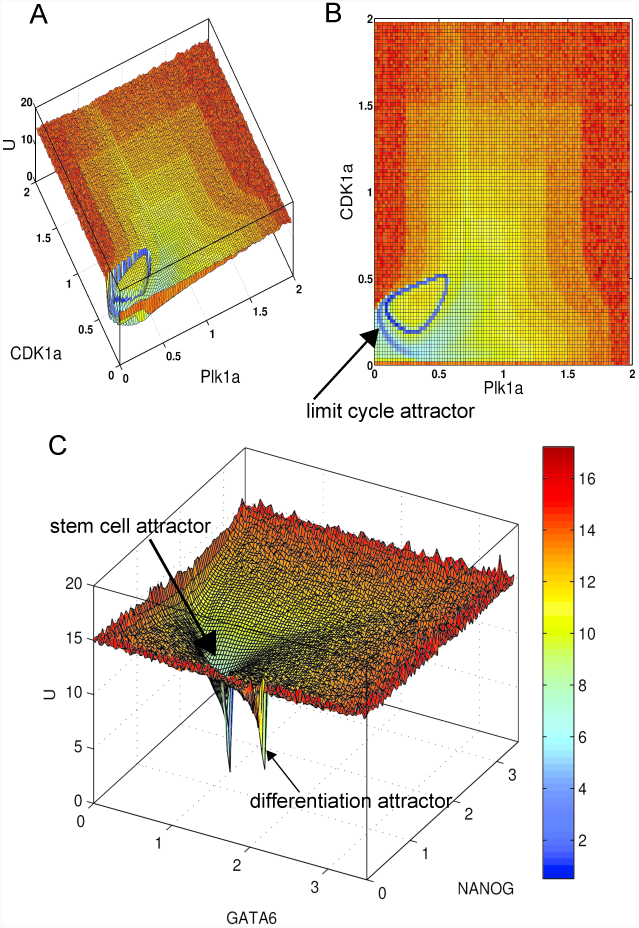
Potential landscapes for Example 4 and Example 5. **(a)** 3D view of the Waddington’s epigenetic landscape for the 3-gene ODE model of cell cycle control which demonstrates the limit cycle attractor (Example 4). **(b)** Top view of the Waddington’s epigenetic landscape for the ODE model of cell cycle control demonstrates the limit cycle oscillator, which corresponds to the limit cycle attractor. **(c)** Waddington’s epigenetic landscape based on a 52-gene network model^13^ (Example 5), which shows two basins of attraction. By comparing the locations of the attractors labeled by Li and Wang^13^, we observe that the stem cell attractor located on the left is the dominant attractor, i.e. bigger and deeper (blue color) than the differentiation attractor located on the right.

The limit cycle is a unique feature of dynamical systems for describing trajectories attracted to a closed-form cycle from inside and outside the closed orbit^23^. It is traditionally illustrated as a closed-form orbit (with no intersection or crossing) in a two-dimensional diagram as shown in the top view of the potential landscape (Figure 3b). Limit cycle oscillations can be viewed as time-course simulations with a fixed periodic form of oscillation. The potential landscape constructed by our method can capture the limit cycle attractor in a 3D view (Figure 3a).

#### Example 5: Stem cell differentiation and reprogramming from Li and Wang^13^

Finally, to test if our method can model high-dimensional gene regulatory networks, we selected a 52-gene network model proposed by Li and Wang^13^ for quantifying the stem cell differentiation and reprogramming. The authors of Ref. 13 identified two attractors in the state space of gene expression: stem cell attractor and differentiated cell attractor. To plot the potential landscape we chose two marker genes GATA6 and NANOG as in Li and Wang^13^, which play pivotal roles in regulating stem cell fates, and used their expression levels as the coordinates of the 2D panel. The shape of our potential landscape is consistent with that in Li and Wang^13^, e.g. both showing two attractors (Figure 3c). The potential landscape shows that the stem cell attractor has a bigger basin of attraction and lower potential value than the differentiated cell attractor.

The result in Figure 3c suggests that our method can capture the transient non-equilibrium states and the attractors of the equilibrium steady states with more details than the results reported by Li and Wang^13^. The potential landscape displays one dominant attractor shown as the stem cell attractor, which implies that the probability of getting attracted to this stem cell attractor is higher than the differentiated cell attractor.

### Computational time

For existing models of Waddington’s epigenetic landscape in the literature, most authors did not report computational time for obtaining potential landscapes. Here, we record the computational time for generating each of the landscape using our Monte Carlo method (Table 1). We used MATLAB R2012b software installed on a Dell desktop computer running Windows 7 (64-bit) operating system with 8 GB memory (RAM). Table 1 shows the benchmark computational time in generating Waddington’s epigenetic landscape. For example, even for the 52-gene model of Li and Wang^13^, our method needs only 33 minutes and 50 seconds. One key factor that might affect the computational time is the non-linearity in the model equations. The 2-gene model from Zhou *et al.*^15^ contains four cubic terms, whereas the 52-gene model from Li and Wang (2013) is composed of Hill functions and as such takes much less time. Moreover, Ferrell *et al.*^19^ model with non-linearity terms of multiplication between one variable and Hill functions required the third longest time although it contains only 3 variables.

**Table 1:** Benchmark computational time for generating Waddington’s epigenetic landscape

| No. | Model | No. variables | No. interactions | Computational time |
| --- | --- | --- | --- | --- |
| 1 | Gardner *et al.* (2000)<sup>17</sup> | 2 | 2 | 10 min 5 sec |
| 2 | Li and Wang (2015)<sup>18</sup> | 6 | 16 | 12 min 0 sec |
| 3 | Zhou *et al.* (2012)<sup>15</sup> | 2 | 8 | 39 min 21 sec |
| 4 | Ferrell *et al.* (2011)<sup>19</sup> | 3 | 3 | 20 min 22 sec |
| 5 | Li and Wang (2013)<sup>13</sup> | 52 | 123 | 33 min 50 sec |

### Landscape modeling based on stochastic Chemical Langevin Equation (CLE)

The models of Waddington’s epigenetic landscape in the previous section were constructed based on deterministic simulations using ODEs. Here, we also investigate the landscape model for Li and Wang^18^ constructed using a stochastic approach based on Chemical Langevin Equation (CLE)^24,25^. We applied the same procedure as described in Algorithm 1 in the Methods Section with the only difference that the numerical simulation of the time-course data was done with CLE instead of ODEs. For the 6-gene network model in Example 2 from Li and Wang^18^, we defined 22 reactions (16 network interactions of activation and inhibition plus 6 self-degradations) and converted the rate constants from the deterministic *k*_*i*_ in ODEs to the stochastic rate constants *c*_*i*_, as explained in Methods. We simulated 100,000 trajectories with random initial conditions. Two examples of the time-course simulation using CLE with random initial conditions are given in Figure 4a. Each trajectory is projected to a 2D plane of dimensions *i* and *j* to estimate the probability distribution of *P* (*x*^*i*^, *x*^*j*^) and the quasi-potential *U* (*x*^*i*^, *x*^*j*^) = −ln *P* (*x*^*i*^, *x*^*j*^) as in the ODE model. A potential landscape for the model of Li and Wang^18^ obtained from CLE is shown in Figure 4b, which is comparable to the one obtained using deterministic ODE-based simulation (Figure 1c). The stochastic noise in the CLE only slightly perturbed the dynamic simulations as the shapes of the attractors in the potential landscape were almost unchanged (even when we used a larger noise than in Higham^25^ original code by changing the volume of the system from *V* = 10^−15^ to *V* = 10^−20^). However, the attractors in the potential landscape from CLE display slightly larger basins of attraction. As a result, two of the attractors that are located close to each other in the ODE-based landscape have been merged into one in the CLE-based landscape (Figure 4b). We also set the volume of the system *V* = 10^−15^ for obtaining a large number of molecules, and the landscapes plotted based on ODE and CLE are almost identical (data not shown). The computational time for generating the potential landscape using the CLE is 57.9 minutes, which is much longer than using the ODEs (12 minutes). This is expected as the stochastic simulation normally takes a longer time due to the high computational cost for simulating all the events of biochemical reactions^26^.

**Figure 4:**
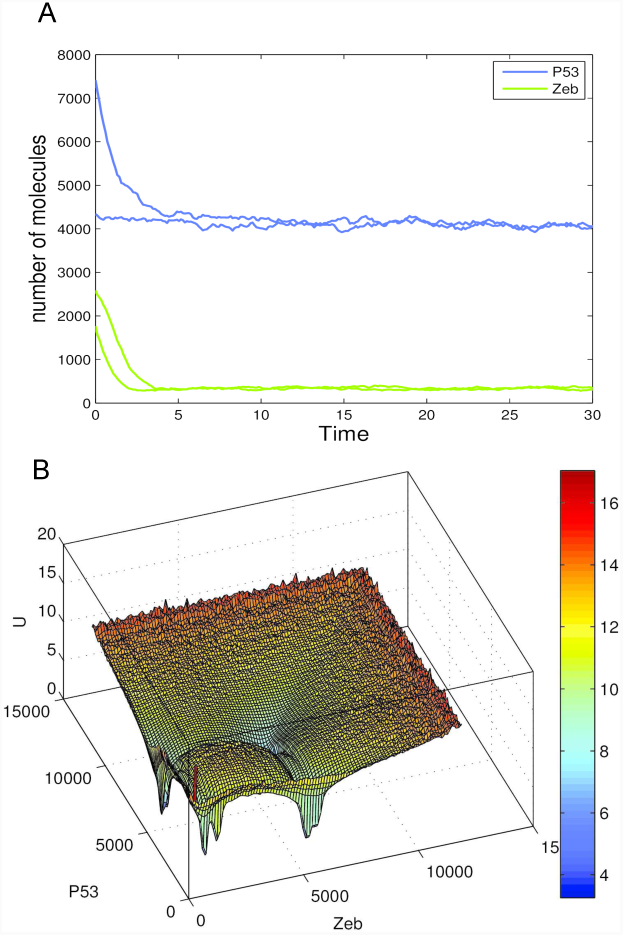
**(a)** Two examples of time-course simulation generated from Chemical Langevin Equation (CLE) with random initial conditions. These time-course simulations show that, starting from two different initial conditions, two trajectories with some randomness can eventually converge to the same attractor. **(b)** Waddington’s epigenetic landscape of the 6-gene network model from Li and Wang^18^, which shows four basins of attraction. The potential landscape was generated by using CLE-based dynamic simulations, and its shape is consistent with that of the landscape obtained using ODEs shown in Figure 1c. It implicates that the attractors in the potential landscape are robust to noise.

We also performed a stochastic simulation using the synthetic toggle switch model of Gardner *et al.*^17^ and obtained similar results as in the Li and Wang^18^ model, i.e. the potential landscape is not affected by the noise (Supplementary Figure S6). These results implicate some degree of robustness of GRNs. The design of the GRN interactions likely enables cells to function properly even though in reality there are intrinsic noises from molecular fluctuations and extrinsic noises from the environment^27,28^. Overall, these results suggest that the global dynamics of the attractors in the Waddington’s epigenetic landscape tends to be robust to noise.

## Discussion

In this paper, we present a novel Monte Carlo method for quantitatively modeling and visualization of Waddington’s epigenetic landscape based on dynamical modeling of GRNs. This method uses a large number of initial conditions randomly sampled from a uniform distribution in the state space, and the collected time-series data are then used to estimate the probability distribution of the cell states and the quasi-potential of the landscape. One key advantage of our method is that it can quantify the potential landscape to display both global and detailed dynamics of the cell state transitions for both equilibrium states (e.g. steady states or attractor states) and nonequilibrium states (e.g. transient states along a kinetic path). The landscape provides a 3D view of the dynamical features (e.g. attractor, saddle point, unstable manifold, stable manifold, limit cycle, and spiral attractor) of the system, whereas conventional dynamical system analysis uses 2D views of bifurcation diagram, phase plane and vector field. Thus, our computational method can be used for detailed modeling and 3D visualization of dynamical systems of cells.

While the method proposed by Li and Wang^13^ formulates the potential as *U* = −ln *P*_*ss*_ (where *P*_*ss*_ is the steady state probability) and uses the self-consistent mean field approximation, our method uses *U* (*x*^*i*^, *x*^*j*^) = −ln *P* (*x*^*i*^, *x*^*j*^) (where *P* (*x*^*i*^, *x*^*j*^) is the estimated probability of an equilibrium steady state or non-equilibrium transient state) and a Monte Carlo method to approximate the potential landscape. Li and Wang^13^ also used 100,000 time-course simulations from random initial conditions in the state space and inferred the steady state probability distribution with multi-variable Gaussian distribution to plot the potential landscape, which displays a smooth surface with basins of attraction. Between their method and ours, the locations of attractors are essentially the same. However, our method can capture more detailed information than their method^13,18^, as demonstrated in Example 2 (with four pairs of attractors as shown in Figure 1c) and in Example 5 (with one dominant attractor as shown in Figure 3c). In Example 2, four pairs of attractors (Figure 1c) can be explained by the kinetic paths for the transition between attractors. As for Example 5, we are not aware of any biological reason for the landscape to display one dominant stem cell attractor and one minor differentiated cell attractor (Figure 3c), which should be investigated in the future. Moreover, our Monte Carlo method is powerful in that it can capture the kinetic path without using the path integral method as in Li and Wang^13^. The kinetic path between two attractors can give biological insight into the transition from one attractor to another attractor where the intermediate state transitions must follow this path towards the final stable state. The kinetic path in the Waddington’s epigenetic landscape can explain why the cell differentiation in embryonic development follows a deterministic path^1,29^ which was called by Waddington himself “chreod”^3,30^.

The potential landscape modeling based on stochastic simulations has also been conducted by Li and Wang (2013) with the method of root mean square distance (RMSD), i.e. using coordinates of two attractors (with locally minimum potentials) as two reference points to reduce a multi-dimensional space into two dimensions of RMSD1 and RMSD2. Using the Langevin dynamics and RMSD they obtained a landscape with the same number of attractors as the landscape based on their self-consistent mean field approximation method. However, the topographies of the two landscapes (from self-consistent mean field approximation and Langevin dynamics with RMSD) reported by Li and Wang^13^ are completely different. Here, we used Chemical Langevin Equation to obtain the time-course trajectories and applied our Monte Carlo method to directly obtain a landscape. Using the stochastic approach of CLE we can obtain a Waddington’s epigenetic landscape consistent with that using the deterministic approach of ODEs. These results of computer simulation highlight the robustness of the gene regulatory network to noise^31,32^.

## Conclusions

The Monte Carlo method for plotting potential landscapes for multi-dimensional GRNs allows us to study the links between genotype and phenotype as initially proposed by Conrad Waddington about 60 years ago^1,2^. Through studies of real biological networks, we have demonstrated the usefulness, simplicity and power of the method for plotting Waddington’s epigenetic landscape. It can facilitate our understanding of cellular differentiation and reprogramming as well as other biological processes. In general, the algorithm proposed here for cellular dynamics can also be applied to studying other types of dynamical systems such as social networks.

## Methods

In this paper, we propose a Monte Carlo method to quantify the potential landscape of equilibrium steady states and non-equilibrium states of a GRN. Firstly, we discuss how to derive the quasi-potential from chemical master equation (CME) using the Monte Carlo method. Instead of solving the CME using the Gillespie’s algorithm (also known as the stochastic simulation algorithm) which is computationally costly, we use numerical simulations of ODE mainly because there are many efficient ODE solvers to obtain time-course trajectories. The Monte Carlo method is used to: (1) generate a large number of random initial conditions that are used for time-course simulations, and (2) estimate probability distribution from time-course trajectories projected into a 2-dimensional plane to quantify the quasi-potential *U*. A summary of the Monte Carlo method for quantifying Waddington’s epigenetic landscape is given in Algorithm 1. Secondly, we discuss how to use stochastic simulations with CLE, which is an improved computation of tau-leaping method for approximate execution of Gillespie’s algorithm^33^. Using the time-course trajectories generated from CLE we can also quantify the Waddington’s epigenetic landscape for a GRN.

### Quantifying Waddington’s epigenetic landscape using deterministic ODE models

#### Derivation of average state probability used in quasi-potential

To derive the quasi-potential for the dynamics of gene expression driving cell state transition, we start from the definition of the CME^34,35^

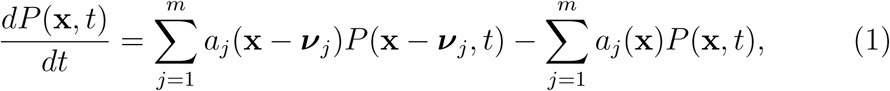

where **x** = (*x*_1_, *x*_2_, *…, x*_*n*_) is the state of the system under study (e.g. a GRN), *x*_*i*_ is the copy number of the *i*th molecular species, *n* is the number of molecular species, *m* is the number of reactions, and *P* (**x**, *t*) is the probability of system in state **x** at time *t*. The function of *a*_*j*_(.) defines the propensity function for the *j*th reaction and ***ν***_*j*_ defines the stoichiometric transition vector for the *j*th reaction. The CME defines the time-evolution of the function *P* (**x**, *t*)^34^. The CME can be interpreted as that the flow of the probability of a system being in state **x** at time *t* is given by the probability of arriving at **x** when reaction *j* fires, *a*_*j*_(**x** *-* ***ν***_*j*_)*P* (**x** *-* ***ν***_*j*_, *t*) subtracted by the probability of the system leaving **x** when reaction *j* fires, *a*_*j*_(**x**)*P* (**x**, *t*)^36^. Summing up all the possible reactions for *j* from 1 to *m* gives Eq. (1). From Eq. (1), we can calculate the probability of a cell in state **x** at time *t*

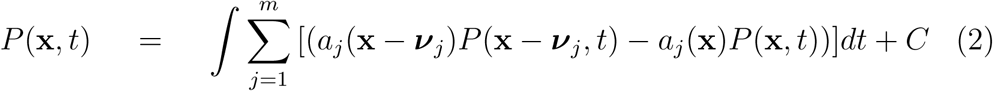

In theory, when the time *t* is large, e.g. approaching infinity, the probability *P* (**x**, **t**) will approach the steady-state probability *P*_*ss*_ ^37,38^. Wang and coworkers^37,38^ proposed a formulation for the quasi-potential as *U* = −ln *P*_*ss*_. However, to study the dynamics of a biological system we also consider the probability over the period of time from 0 to *T*, and calculate the average probability of state at **x** over the time from 0 to *T*, as

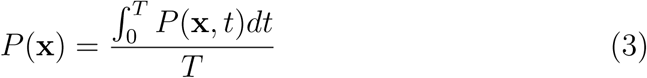

According to the definition of the quasi-potential *U* proposed by Wang and co-workers as *U* = −ln *P*(**x**)^31,38^, thus we approximate the quasi-potential *U* by using *P*(**x**) from Eq.(3) as given by

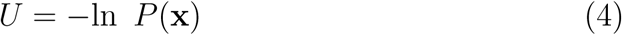

### Using Monte Carlo simulation and Gillespie’s algorithm to estimate quasi-potential

It is difficult to solve *U* = −ln *P*(**x**) analytically. Thus, we can use a Monte Carlo method to get an approximate solution. Monte Carlo methods use computer simulations with random sampling to approximate the exact solutions^39^. It has been widely used for solving a variety of problems including landscape modeling. For example, two recent works used Monte Carlo simulations to model epigenetic landscape and cell type transitions. Nakagawa and Narikiyo^40^ proposed a minimal modeling of epigenetic landscape based on the fitness of interacting cells. Wang *et al.*^41^ proposed a Monte Carlo method based on an ensemble of parameters to simulate the global dynamics of the epigenetic state network. Our Monte Carlo method is different from the above two methods, in that we use a large number of random initial conditions for simulating trajectories and then obtain the probability distribution of *P*(**x**).

First, from Eq. (3) we discretize the formulation from integral into summation:

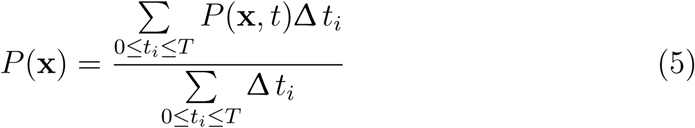

where Δ *t*_*i*_ is the increment of time for *P* (**x**, *t*). Suppose the initial condition is uniformly distributed, and *x*_*i*_ is not larger than *X*_*i*_, which is a positive integer fixed by a modeler based on the maximum value in the dynamical system. Then the probability averaged over the initial conditions is given by:

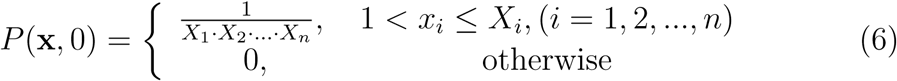

Next, we randomly choose *N* initial states {**x**_1_(0), **x**_2_(0), *….*, **x**_*N*_(0)} which are uniformly distributed in every dimension. Based on these random initial conditions we simulate *N* trajectories using Gillespie’s algorithm^34^. Let us denote the *i* th trajectory by 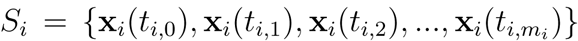, where 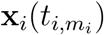 is the state at time 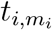 for the *i* th trajectory. From these trajectories we can estimate the probability in Eq. (5) as follows:

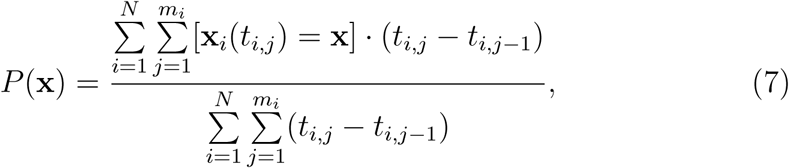

where [*x*] is the Iverson bracket defined by

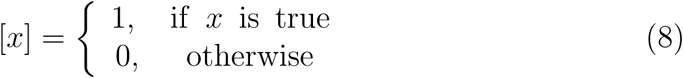

From this Monte Carlo method we can estimate the probability distribution of the state at **x** given by *P*(**x**) and then using Eq. (4) we can obtain the quasi-potential *U* for the dynamical system.

### Improving the speed for obtaining quasi-potential by using ODEs

Using the Gillespie’s algorithm we can simulate the time-evolution trajectories for CME. However, it incurs very high computational cost for simulating every event of the chemical reactions^26^. According to the Gillespie’s work^34^, when the number of molecules present in the biochemical reactions is large the stochastic and deterministic simulation results are almost equivalent with negligible random fluctuations. Assuming that the number of molecules present in the biochemical reactions is large, we use numerical solution of ODEs to speed up the computation. We also randomly choose *N* initial conditions {**x**_1_(0), **x**_2_(0), *…*, **x**_*N*_(0)} which are uniformly distributed in every dimension. However, these initial conditions are measured in concentration levels of the molecular species. Using these random initial conditions we simulate *N* trajectories.

After numerically solving the ODEs, the output trajectories can be discretized from the continuous time with a specific time step. As such, we can use Eq. (7) to obtain *P*(**x**) for calculating the quasi-potential of the dynamical system. The only difference here is that **x** is measured in concentration of each molecular species instead of the number of molecules as in the Gillespie’s algorithm. We choose to use an ODE solver with a fixed time step so that the time difference for Δ*t* is constant. Let us assume the time difference is given by

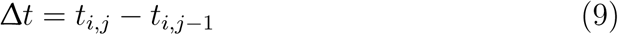

Then, substituting Eq. (9) into Eq. (7) and making simplification, we obtain

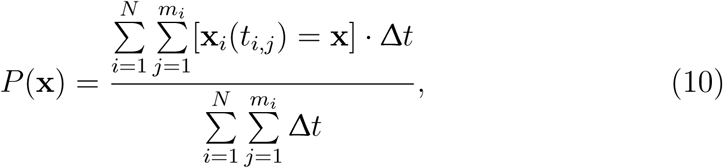

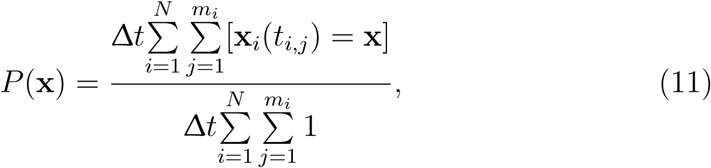

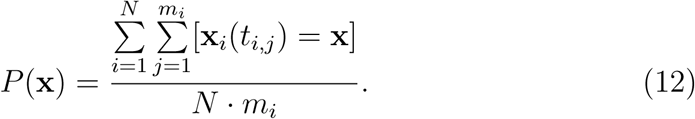

With Eq. (12) we have simplified Eq. (7) and improved the calculation speed to obtain the quasi-potential value. However, accurate estimation of *P*(**x**) still involves high computational cost due to the large state space of the system. In order to further improve the speed of the computation we apply a coarse graining formulation defined by

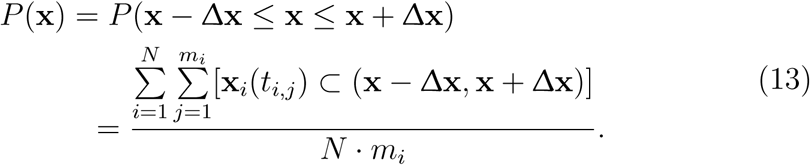

The coarse graining formulation above is implemented as the division of a 2-D plane into grid boxes.

### Plotting landscape

From previous sections we have derived Eq. (7) and Eq. (13) for calculating *P*(**x**). However, these equations are for dynamical systems with high-dimensional state spaces. In order to plot the potential landscape in 3 dimensions for human viewers to understand, we need to reduce the dimensions of a system. There are many dimensionality reduction methods that can be applied to the plotting of landscape. Here, we propose a simple method of dimensionality reduction by projecting the trajectories into a 2-dimensional plane. By marginalizing out all the variables except for the *i*th and *j*th variables, we can obtain the probability distribution of the states in the *i*th and *j*th dimensions as

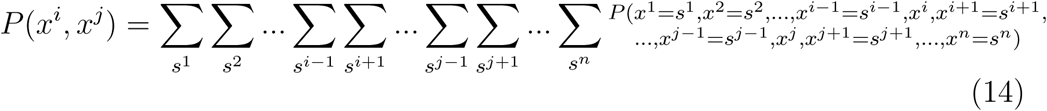

After the dimensionality reduction, we can subtitute Eq. (14) into Eq. (4) to obtain the quasi-potential of a state in two dimensions

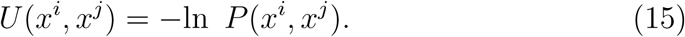

With *x*^*i*^, *x*^*j*^ and *U* (*x*^*i*^, *x*^*j*^) as the *x, y* and *z* axes respectively, we can plot the landscape in 3 dimensions. Since the calculation of *P* (*x*^*i*^, *x*^*j*^) considers the time course from *t*_0_ to 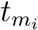, our method can be used to analyze the properties of transient states, rather than being limited to steady states, and thereby can reveal more dynamical details.

Our Monte Carlo method is summarized in Algorithm 1. Essentially, the algorithm collects *N* simulated time-course trajectories from random initial conditions in the state space. Here, we use a fixed time step of Δ*t* = 0.1 for numerical simulation, and thereby we can use the number of time points along a trajectory instead of the length of continuous time to estimate the probability of a state. Then from these time-course data, we project the points of the trajectories into a phase plane of two selected variables of interest and estimate the probability. For example, Figure 5 shows how the probability distribution can be estimated in a plane that has been divided into grid boxes. A grid box with a locally maximal number of points on the trajectories represents an attractor. In Figure 5 there are two yellow grid boxes with locally maximal numbers of points from the trajectories, and therefore they represent two attractors. These results are then used to estimate the quasipotential *U* (*x*^*i*^, *x*^*j*^) = *-*ln *P* (*x*^*i*^, *x*^*j*^), where *P* (*x*^*i*^, *x*^*j*^) is the probability of a state which is either an equilibrium state or a non-equilibrium state. A non-equilibrium state quantifies the transient behaviors of the system such as the intermediate flows in a vector field, whereas an equilibrium state quantifies a repeller (unstable steady state) or an attractor (stable steady state)^42^. This formulation and approximate calculation of the quasi-potential enables us to plot the Waddington’s epigenetic landscape with details.

**Figure 5:**
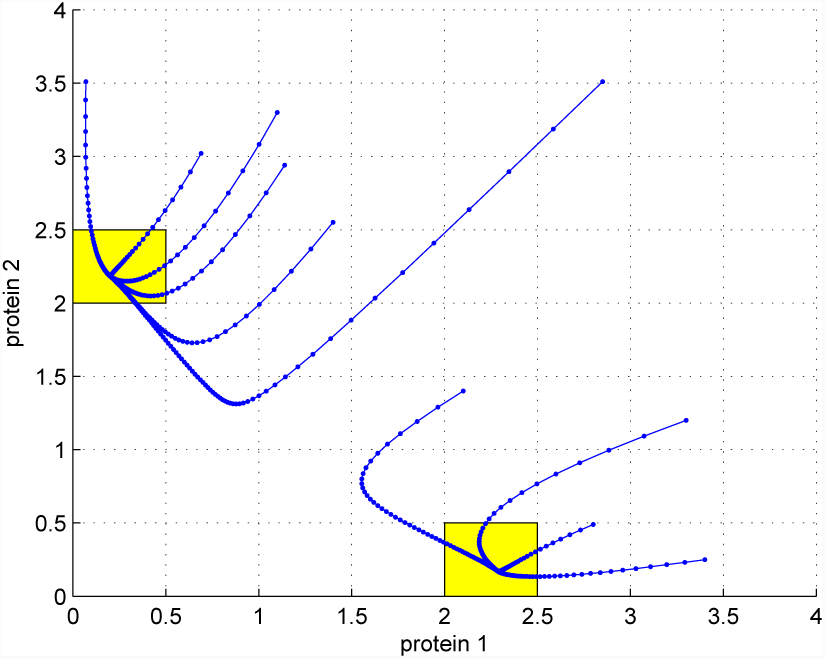
Illustration of the Monte Carlo method for approximating the probability distribution and identifying attractors. The projection of the time-course data into a plane with grid boxes enables the estimation of the probabilities of cellular states. In this example 10 trajectories (blue lines) are shown and the plane is split into 8 × 8 grid boxes. A grid box with a locally maximal number of points of trajectories corresponds to an attractor. In this landscape, there are two attractors as indicated by the two yellow boxes.

#### Algorithm 1.

Monte Carlo steps for quantifying quasi-potential landscape

1. generate *N* random initial conditions
2. generate 2-dimensional grids
3. for each initial condition:
  i. generate one trajectory
  ii. project the trajectory to the 2-dimensional grid boxes
  iii. from the trajectory calculate the number of points in each grid box
4. estimate probability of each grid box *P* (*x*^*i*^, *x*^*j*^) as given by Eq. (15)
5. calculate quasi-potential of each grid box using *U* (*x*^*i*^, *x*^*j*^) = *-*ln *P* (*x*^*i*^, *x*^*j*^)
6. plot the landscape in 3 dimensions

### Stochastic modeling of landscape based on Chemical Langevin Equation (CLE)

To test the role of noise in gene expression and the robustness of GRNs we also investigate a second type of potential landscape that is based on stochastic time-course simulations. The algorithm for obtaining the potential landscape is similar to Algorithm 1. The key difference is that the stochastic approach uses Chemical Langevin Equation (CLE)^24^, an improved version of the tau-leaping method for approximation of the Gillespie’s algorithm^33^, in simulating the time-series trajectories. CLE differs from ODEs in that the biochemical reactions are simulated using the stochastic part in the CLE^25^. We adapted the CLE code from Higham^25^ with the model reactions from the GRNs of Li and Wang^18^ and Gardner *et al.*^17^. We used a larger noise than in the original code of Higham^25^, by decreasing the volume of the system from *V* = 10^−15^ to *V* = 10^−20^. Since CLE is a well-established method for stochastic simulation of biochemical reactions^33^ we will not discuss it in detail here. The CLE and their implementation in MATLAB code are given in Supplementary material. Below we will illustrate how to construct CLE by converting from ODEs. The definitions of the rates of change of molecular species are different between ODEs and CLE. In ODEs the molecular species are measured in concentrations, whereas in CLE the molecular species are measured in the numbers of molecules. To explain how to convert the deterministic rate constant *k*_*i*_ in ODEs to the stochastic reaction rate constant *c*_*i*_ in CLE, let us look at Eq. (16), an example ODE model equation for protein *x* with self-activation (the first term on the right hand side) and spontaneous degradation (the second term):

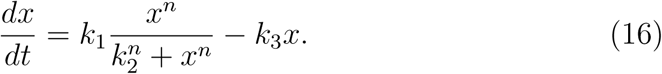

Let us use *X* to denote the copy number of protein *x*. Hereafter we will use *x* to denote the concentration of protein *X*, in the unit of *μM*. The relationship between *x* and *X* is given by:

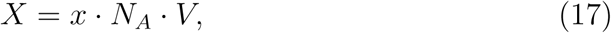

where *N*_*A*_ is the Avogadro’s number which equals 6.023 × 10^23^ and *V* is the volume of the system in liters^25^. Next, denote *B* = *N*_*A*_ · *V*, then Eq. (17) becomes

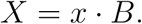

Multiply *B* to both sides of Eq. (16), and we get

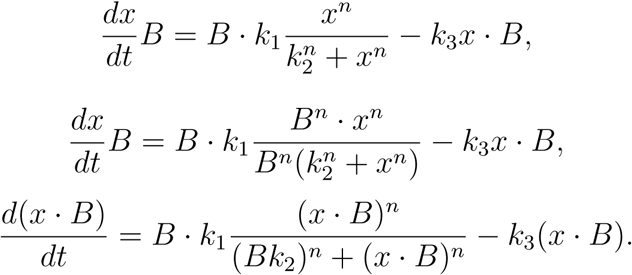

Since *X* = *x · B*, we obtain the following equation:

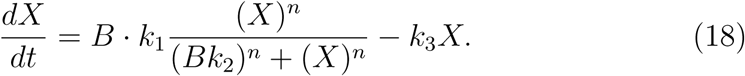

In the stochastic approach the rate of change for *X* is defined by:

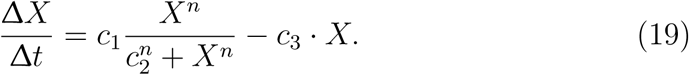

Comparing Eq. (18) and Eq. (19), we deduce that,

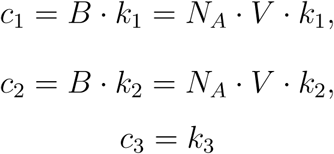

From the derivations above, we demonstrate the relationship between *k*_*i*_ and *c*_*i*_ which enables us to obtain the *c*_*i*_ for the stochastic simulation using CLE.

## Supporting information

User manual for running our source code

MATLAB source code of our Monte Carlo method and case study models

## Availability of data and materials

All the data and materials are provided in the additional files.

## Author’s contributions

XZ and KHC created the method and implemented the algorithm and case studies. JZ supervised the design of the method and the study. XZ, KHC and JZ wrote the manuscript.

## Acknowledgements

The authors would like to thank Chunhe Li and Sudin Bhattacharya for answering our questions regarding their published methods. We also thank Bard Ermentrout for the permission to use the XPPAUT software for plotting the phase plane in Figure 2c.

