## Supplementary material for "A Monte Carlo method for *in silico* modeling and visualization of Waddington’s epigenetic landscape with intermediate details": User manual for running our source code

### 1 Drawing Waddington's epigenetic landscape using ODEs

#### 1.1 Introduction

Our algorithm for quantifying Waddington's epigenetic landscape was implemented in MATLAB. The source codes are distributed in different folders for different models of gene regulatory network (GRN) (as discussed in our paper). This document explains how you can run the MATLAB code to plot the Waddington's epigenetic landscape.

This MATLAB code package in the folder named "ODEs" can be modified according to user's own GRN model in ordinary differential equations (ODEs), which can be encoded in the `equations.m` file. For details please read Section 1.4 titled "Using the code to plot potential landscapes for your gene networks in ODEs."

#### 1.2 Usage

For different GRNs the model equations are different, so there are 5 folders for the 5 models presented in the main text. Open a model folder and you can run the MATLAB code to plot the Waddington's epigenetic landscape. Each folder contains 2 MATLAB files:

1. `Setting_and_running.m`
2. `equations.m`

Five common MATLAB files are located at a common code folder.

1. `GetAllInitialConditions.m`
2. `GetAllTrajectories.m`
3. `GetPositionProbabilities.m`
4. `GetOneTrajectory.m`
5. `DrawLandscape.m`

To run and plot the Waddington's epigenetic landscape, open the `Setting_and_running.m` file in MATLAB editor and click the Run button in the menu bar (see Fig. S1) or press F5.

Alternatively, you can run and plot the landscape in the MATLAB Command Window. Type the filename and then press Enter at the command line as below:

```
>>Setting_and_running
```

After the calculation is finished with $N$ trajectories ($N$ is set to 100,000 by default but user can set $N$ to a different integer number that is nonnegative in the file `Setting_and_running.m`), you will obtain two graphs: the 3D view and top view of the landscape.

#### 1.3 Examples

**Example 1: Bistable synthetic toggle switch from [1]**

The GRN is given in Fig. S2. According to the Hill function formulation, the two equations for the GRN are:

$$\frac{du}{dt} = \frac{\alpha_1 \cdot S^3}{S^3 + v^3} - k_1 u, \tag{S1}$$

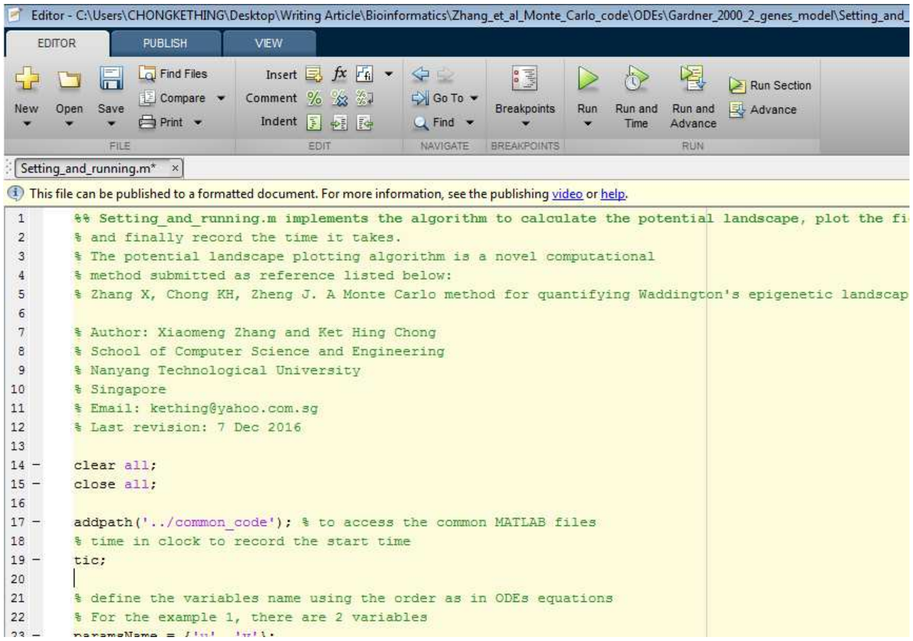


**Figure S1:** Running the MATLAB code using Editor. Click the Run button (green arrow) or by pressing F5 function key.

$$\frac{dv}{dt} = \frac{\alpha_2 \cdot S^3}{S^3 + u^3} - k_2 v, \tag{S2}$$

We used the model parameters given in [2, 3] where $\alpha_1 = 3$, $\alpha_2 = 3$, $S = 1$, $k_1 = 1$ and $k_2 = 1$.

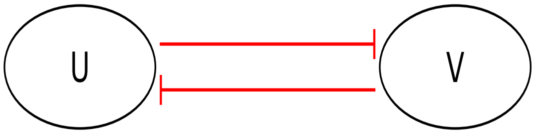


**Figure S2:** The network interactions for the model in Example 1, which is a simple synthetic toggle switch that contains two genes $u$ and $v$. The two genes inhibit each other.

To plot the landscape, open the folder `Gardner_2000_2_genes_model`. Type the filename at the command line as below:

```
>>Setting_and_running
```

After the calculation completes with $N = 100{,}000$ trajectories, the 3D view and top view of the Waddington's epigenetic landscape will be plotted. Finally, the time taken for the calculation and plotting of the landscape will be printed in the command window. The potential landscape displays 2 attractors as shown in Fig. S3.

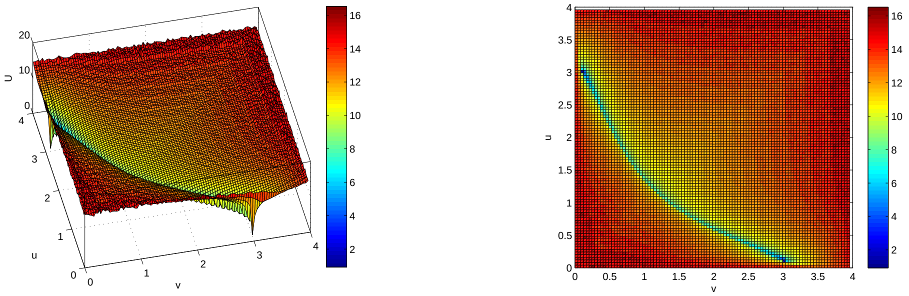


**(a)** 3D View

**(b)** Top View

**Figure S3:** Plots of the landscape of the bistable synthetic toggle switch from [1].

**Example 2: Quantification of Waddington's epigenetic landscape of the GRN model of cancer attractors** [4]

The model of GRN from Li and Wang (2015) [4] is shown in Fig. S4. We used model equations given in [4] as listed below:

$$\frac{dP53}{dt} = \frac{sa \cdot P53^4}{S^4 + P53^4} + \frac{b \cdot S^4}{S^4 + MDM2^4} - k \cdot P53 \qquad (S3)$$

$$\frac{dMDM2}{dt} = \frac{a \cdot P53^4}{S^4 + P53^4} + \frac{b \cdot S^4}{S^4 + (miR145)^4} - k \cdot MDM2 \qquad (S4)$$

$$\frac{dOCT4}{dt} = \frac{sa_1 \cdot (OCT4)^4}{S^4 + (OCT4)^4} + \frac{b \cdot S^4}{S^4 + P53^4} + \frac{b \cdot S^4}{S^4 + (miR145)^4} - k \cdot OCT4 \qquad (S5)$$

$$\frac{d(miR145)}{dt} = \frac{a \cdot P53^4}{S^4 + P53^4} + \frac{b \cdot S^4}{S^4 + (OCT4)^4} + \frac{b \cdot S^4}{S^4 + ZEB^4} - k \cdot miR145 \qquad (S6)$$

$$\frac{d(ZEB)}{dt} = \frac{sa_2 \cdot ZEB^4}{S^4 + ZEB^4} + \frac{b \cdot S^4}{S^4 + (miR145)^4} + \frac{b \cdot S^4}{S^4 + (miR200)^4} - k \cdot ZEB \qquad (S7)$$

$$\frac{d(miR200)}{dt} = \frac{a \cdot P53^4}{S^4 + P53^4} + \frac{a \cdot (OCT)^4}{S^4 + (OCT)^4} + \frac{b \cdot S^4}{S^4 + (ZEB)^4} - k \cdot (miR200) \qquad (S8)$$

where $a = 0.5$, $sa = 0.8$, $sa_1 = 0.8$, $sa_2 = 0.8$, $b = 0.1$ , $S = 0.5$ and $k = 1$.

Open the folder `Li_2015_6_genes_model`, and type the filename at the command line as below:

```
>>Setting_and_running
```

After the calculation finishes, the 3D and top views of the Waddington's epigenetic landscape will be plotted. The potential landscape is shown to have four pairs of attractors as shown in Fig. S5.

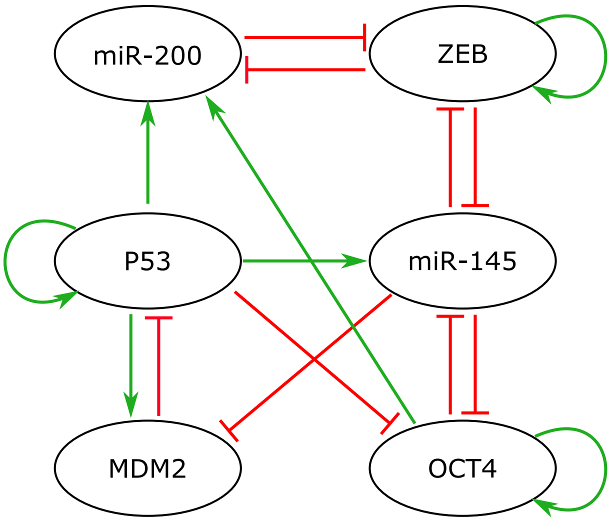


**Figure S4:** The GRN model of Example 2 from [4] contains 7 activations and 9 inhibitions, which are represented by green arrows and red lines with a short bar respectively.

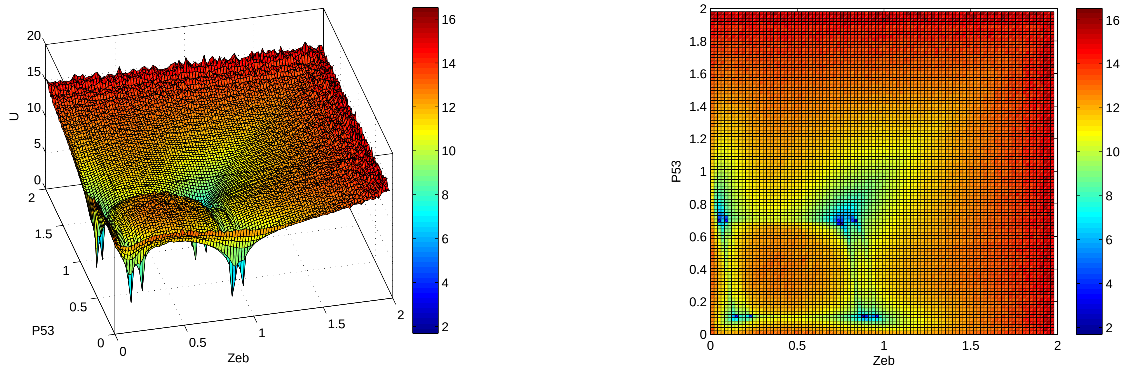


**(a)** 3D View

**(b)** Top View

**Figure S5:** Plots of Waddington's epigenetic landscape of the GRN model of Li and Wang (2015) [4].

##### 1.4 Using the code to plot potential landscapes for your gene networks in ODEs

In order to plot a potential landscape of your own model, you only need to edit two files: `equations.m` and `Setting_and_running.m`.

First, define your gene network model in ODEs in the file of `equations.m`. Secondly, change the model setting in the file of `Setting_and_running.m`. Set the variable names (`variableNames`) as the order of the equations according to the ODEs (e.g. for the Example 1, the `variableNames = {'u', 'v'}`). Then, you need to set a different maximum range of the state space for your gene network. For example, based on the prior knowledge from time-course simulations the maximum value of the level of a protein is say 3, then you may enter the maximum value as 4 (`range_max` = 4). We added a margin of 1 such that the landscape will cover enough values for getting a complete landscape. Because each GRN model equations and landscape is different, user may need to adjust and add a suitable margin value.

In order to set the state space for the random initial conditions, change the dimension of your model equations in ODEs. For example, if your ODEs contain $n$ variables, then set:
`initialRange = [zeros(1,n)+range_min; zeros(1,n)+range_max];`

Also, you need to change the `index = [variable_1 variable_2]` to the two indices of genes (or proteins) of interest to you. If your model equations contain only 2 variables, then set the index to:
`index = [1 2]` where `variable_1` = 1 and `variable_2` = 2.

Finally, you can run the calculation to plot the Waddington's epigenetic landscape as in the two aforementioned examples.

#### 2 Drawing Waddington's epigenetic landscape using stochastic method of CLE

##### 2.1 Introduction

This section explains how to run the MATLAB code to generate the Waddington's epigenetic landscape using a stochastic approach with Chemical Langevin Equation (CLE). The CLE implementation was adapted from [5].

This MATLAB code package in the folder named "CLE" can be modified according to your GRN model in ODEs. The ODEs can be converted into CLE with different rate constants and stoichiometric matrix $\boldsymbol{V}$ which should be defined in the `GetTrajectory.m` file. The definition of CLE and the explanation of how to convert ODEs into CLE are given in Section 2.4. For details read Section 2.4 titled "Using the code to plot potentials landscape for your gene networks in CLE. "

##### 2.2 Usage

Open the folder of the model that you wish to run to plot the Waddington's epigenetic landscape. In our package, we have provided files for two models, Example 1 and Example 2 which correspond to the two examples of ODE model in the preceding section. Each folder contains 2 MATLAB files:

1. `Setting_and_running_CLE.m`
2. `GetTrajectory.m`

Three common MATLAB files are located at a common code folder for CLE.
1. `GetPositionProbabilities.m`

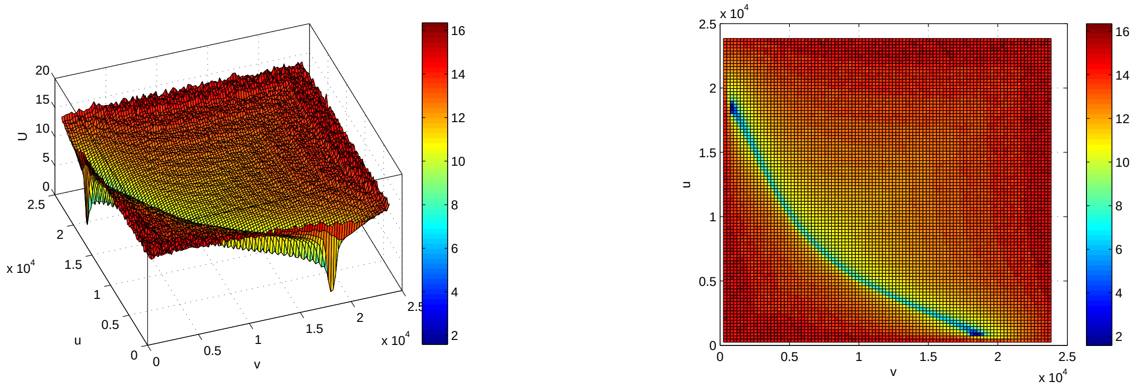


**(a)** 3D View **(b)** Top View

**Figure S6:** Plots of the landscape for the bistable synthetic toggle switch from [1] using CLE.

2. `GetAllTrajectories.m`
3. `DrawLandscape.m`

To plot the Waddington's epigenetic landscape, open the first m file `Setting_and_running_CLE.m` in MATLAB editor and click the Run button on the menu bar.

After the calculation finishes with $N$ simulated trajectories, user should obtain two graphs: the 3D view and the top view of the landscape. The default value of $N$ is 100,000 but user can modify the code to set the value of $N$.

#### 2.3 Examples

**Example 1: Bistable synthetic toggle switch from [1]**

We use the ODE model equations given in Eqn (S1) and Eqn (S2) [2].

Open the folder `Gardner_2000`. Type the filename at the command line as below:

```
>>Setting_and_running_CLE
```

After the calculation finishes with $N$ = 100,000 simulated trajectories, the 3D and top views of the Waddington's epigenetic landscape will be plotted. Moreover, the time taken for the calculation and plotting of the landscape will be displayed in the command line.

In the potential landscape there are two attractors (see Fig. S6), which is consistent with the ODE-based landscape in Fig. S3.

**Example 2: Cancer attractor landscape of from [4]**

We use the ODE model equations given in Eqn (S3-S8) [4]. Open the folder `Li_2015_6_genes_model`. Type the filename at the command line as below:

```
>>Setting_and_running_CLE
```

After the calculation finishes, the 3D and top views of the Waddington's epigenetic landscape will

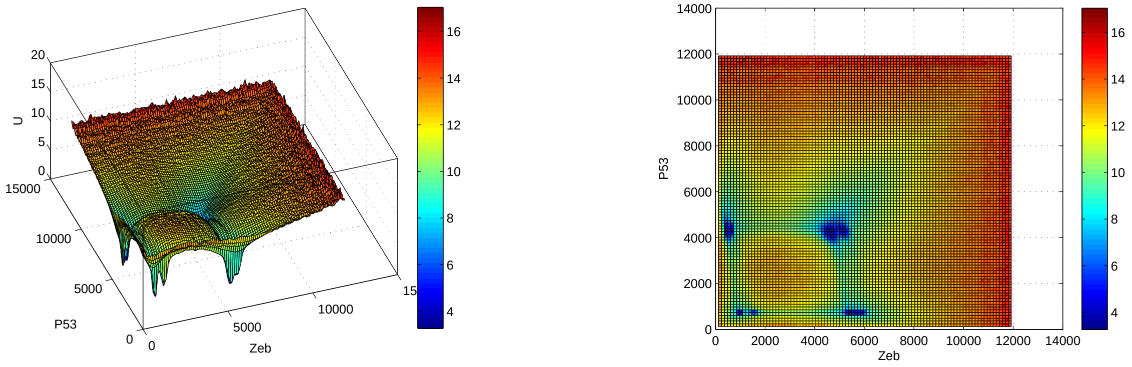


**(a)** 3D View **(b)** Top View

**Figure S7:** Plots of Waddington's epigenetic landscape of the GRN model in [4] using CLE.

be plotted. The resulting potential landscape (see Fig. S7) will display four pairs of attractors which is consistent with that in Fig. S5.

#### 2.4 Using the code to plot potential landscapes for your gene networks in CLE

In order to plot the potential landscape of your model, you need modify two files: `GetTrajectory.m` and `Setting_and_running_CLE.m`. For a basic understanding of CLE, readers are referred to a guide with MATLAB code given by Higham [5]. Here, we list the formula for CLE and discuss how to implement the model from [1]. The formula for CLE as given in [5] is:

$$Y(t+\tau) = Y(t) + \tau \sum_{j=1}^{m} \nu_j a_j(Y(t)) + \sqrt{\tau} \sum_{j=1}^{m} \nu_j \sqrt{a_j(Y(t))} Z_j \tag{S9}$$

where $Y(t)$ is the state vector, $\tau$ is the leap time, and $\nu_j$ is the state-change vector for the $j$th type of reaction, for $1 \leq j \leq m$, $\nu_j \in \mathbb{R}^n$ for $m$ reactions and $n$ molecular species. $\boldsymbol{V}$ is the stoichiometric matrix, which is an $n$ x $m$ matrix, where $\nu_j$ is the $j$th column. $a_j(Y(t))$ is the propensity function for the $j$th reaction, which means the probability of the $j$th reaction occurring in the infinitesimal time interval $[t, t+\tau)$ is given by $a_j(Y(t))\tau$. $Z_j$ is a random number generated from the uniform distribution.

##### 2.4.1 Defining model equations into reactions

First define your gene network model reactions in CLE in the file `GetTrajectory.m`. A model in ODEs can be converted into CLE with the different rate constants $c_i$ from the ODEs rate constants $k_i$ and stoichiometric matrix $\boldsymbol{V}$.

For instance, let us look at the model with ODEs from [1] given below:

$$\frac{du}{dt} = k_1 \frac{k_2^3}{(k_2^3 + v^3)} - k_3 u \tag{S10}$$

$$\frac{dv}{dt} = k_4 \frac{k_2^3}{(k_2^3 + u^3)} - k_5 v \tag{S11}$$

In the first ODE, it contains two reactions: the inhibition by $v$ and the self-degradation of $u$. Similarly the second ODE also contains two reactions. Correspondingly, there are 4 reactions (R1-R4) and 5 rate constants: $k_1$, $k_2$, $k_3$, $k_4$ and $k_5$.

$$R1 =: k_1 \frac{k_2^3}{(k_2^3 + v^3)},$$

$$R2 =: -k_3 u,$$

$$R3 =: k_4 \frac{k_2^3}{(k_2^3 + u^3)},$$

$$R4 =: -k_5 v.$$

From the formulas of R1-R4, we can determine the stoichiometric matrix $\boldsymbol{V}$ and the stochastic rate constants.

##### 2.4.2 Defining stoichiometric matrix

The stoichiometric matrix $\boldsymbol{V}$ for this example contains 4 columns, each representing one reaction. User can key in column by column of the matrix $\boldsymbol{V}$. From the ODEs in Eqn (S10) and Eqn (S11) the term with a positive rate means synthesis of protein and a negative rate means degradation of protein. If a reaction is positive put 1 (meaning that the number of molecule is increased by one), and if the reaction is negative put $-1$ (meaning the number of molecule is reduced by one). In this example, each row represents one molecular species where row 1 represents $u$ and row 2 represents $v$.

R1: $u$ increases by 1, first column $\nu_1 = \begin{pmatrix} 1 \\ 0 \end{pmatrix}$;

R2: $u$ decreases by 1, second column $\nu_2 = \begin{pmatrix} -1 \\ 0 \end{pmatrix}$;

R3: $v$ increases by 1, third column $\nu_3 = \begin{pmatrix} 0 \\ 1 \end{pmatrix}$;

R4: $v$ decreases by 1, fourth column $\nu_4 = \begin{pmatrix} 0 \\ -1 \end{pmatrix}$.

Thus, the stoichiometric matrix $\boldsymbol{V}$ is given by:

$$\boldsymbol{V} = \begin{pmatrix} 1 & -1 & 0 & 0 \\ 0 & 0 & 1 & -1 \end{pmatrix}.$$

##### 2.4.3 Defining stochastic rate constants $c_i$ in CLE

Based on our paper Methods in Section 4.2 we can determine the relationship between deterministic rate constants $k_i$ and stochastic reaction rate constants $c_i$. The stochastic reaction rate constants $c_i$ are:

$c_1 = N_A \cdot V \cdot k_1$
$c_2 = N_A \cdot V \cdot k_2$
$c_3 = k_3$
$c_4 = N_A \cdot V \cdot k_4$
$c_5 = k_5$

where $N_A$ is the Avogadro's number $N_A = 6.023 \times 10^{23}$ and $V$ is the volume of the system.

Once we know how to convert the rate constants from $k_i$ to $c_i$ we can use $c_i$ to implement into MATLAB code. Here, we illustrate the implementation of it.
In ODEs rate constants *($k_i$)* are:

```
k1 = 3;
k2 = 1;
k3 = 1;
k4 = 3;
k5 = 1;
```

Thus, in CLE rate constants *($c_i$)* are:

```
nA = 6.023e23; % Avogadro's number
vol = 1e-20; % system's volume
c(1) = 3*(nA*vol);
c(2) = 1*nA*vol;
c(3) = 1;
c(4) = 3*(nA*vol);
c(5) = 1;
```

In Higham's paper [5], the volume of the system is vol=$10^{-15}$. To mimic a higher level of noise, however, we use a smaller value of volume by setting vol = $10^{-20}$ and obtained consistent landscape (Fig. S7).

##### 2.4.4 Defining *a(j)* the propensity function

The propensity function for the $j$th reaction is given by $a_j(Y(t))$. Thus, the expressions of $a(j)$ are given by:

$$a(1) = \frac{c(1)*c(2)^3}{c(2)^3+Y(2)^3},$$

$$a(2) = c(3)*Y(1),$$

$$a(3) = \frac{c(4)*c(2)^3}{c(2)^3+Y(1)^3},$$

$$a(4) = c(5)*Y(2).$$

##### 2.4.5 Implementing the CLE

When all the model reactions and rate constants have been defined, the CLE formula given in Eqn (S9) above is translated into MATLAB code (particularly the line of code highlighted in yellow) as below:

```
tfinal = 30; % simulation time
L = 250; % stepsize
tau = tfinal/L;
for k=1:L
    a(1) = c(1)*c(2)^3/(c(2)^3+Y(2)^3);
    a(2) = c(3)*Y(1);
    a(3) = c(4)*c(2)^3/(c(2)^3+Y(1)^3);
    a(4) = c(5)*Y(2);
    d(1) = tau*a(1) + sqrt(abs(tau*a(1)))*randn;
```

```
    d(2) = tau*a(2) + sqrt(abs(tau*a(2)))*randn;
    d(3) = tau*a(3) + sqrt(abs(tau*a(3)))*randn;
    d(4) = tau*a(4) + sqrt(abs(tau*a(4)))*randn;
    %  Y(t+tau)= Y(t) + tau*(aj*vj) + sqrt(tau*aj)*Zj*vj
     Y = Y + d(1)*V(:,1) + d(2)*V(:,2) + d(3)*V(:,3)+ d(4)*V(:,4);
    Yvals(:,k+1) = Y; % store all the state values into matrix Yvals
end
```

##### 2.4.6 Initializing the model setting

Finally, modify the model setting in the file of `Setting_and_running_CLE.m`. Set the variable names (`variableNames`) as the order of the variables (or proteins) according to the CLE in the stoichiometric matrix. Then, you need set a different range of the state space for your gene network. For example, based on the prior knowledge from ODEs time-course simulations the maximum value of the protein level is say 3, then you may set the range to 4 (`range_max` = 4).
You need to change the dimension of your model in CLE. For example, if your CLE contain 2 variables, then set:

$$\begin{aligned} initialRange = [&zeros(1,2) + \texttt{range\_min}; \%\ set\ the\ minimum\ range \\ &zeros(1,2) + \texttt{range\_max}]; \%\ set\ the\ maximum\ range \end{aligned}$$

Also, you may change the index [`variable_1` `variable_2`] as the two genes or proteins of interest to you. If your model equations only contain 2 variables, then `variable_1` = 1 and `variable_2` = 2. Then, you can run the calculation to plot the Waddington's epigenetic landscape as given in the example above.

### 3 Contact information

The MATLAB codes were written by Xiaomeng Zhang and Ket Hing Chong.

For any question or comments, please contact Jie Zheng <>
