## Supplementary material for "A Monte Carlo method for *in silico* modeling and visualization of Waddington’s epigenetic landscape with intermediate details": MATLAB source code of our Monte Carlo method and case study models: DrawLandscape.m

```
% Get all time course simulations using CLE
allTrajectories=GetAllTrajectories(initialConditions, vol);

%% Calculate probability distribution


positionProbabilities = GetPositionProbabilities(fromInitialCondition*nA*vol, toInitialCondition*nA*vol, index, splitNumber, allTrajectories);

% Normalized Probability
normalized_Prob=positionProbabilities./(sum(sum(positionProbabilities)));

% U = - ln P
Neg_lnP=-log(normalized_Prob);

[x y] = meshgrid(linspace(fromInitialCondition(index(1))*nA*vol, toInitialCondition(index(1))*nA*vol, splitNumber), ...
    linspace(fromInitialCondition(index(2))*nA*vol, toInitialCondition(index(2))*nA*vol, splitNumber));

%% Plotting the potential landscape
% 3D view
figure;
surf(x,y,Neg_lnP);
xlabel(variableNames(index(2)));
ylabel(variableNames(index(1)));
zlabel('U')

view(-20,54); 
box on;
colorbar;

%% 
% Top view
figure;

surf(x,y,Neg_lnP);
xlabel(variableNames(index(2)));
ylabel(variableNames(index(1)));
zlabel('U')

view(0,90); 
box on;
colorbar;

disp(['time taken to compute the landscape is ' num2str(toc/60) ' min.'])
```
