## Supplementary material for "A Monte Carlo method for *in silico* modeling and visualization of Waddington’s epigenetic landscape with intermediate details": MATLAB source code of our Monte Carlo method and case study models: GetPositionProbabilities.m

```
function res = GetPositionProbabilities(fromInitialCondition, toInitialCondition, index, splitNumber, allTrajectories)
% This function count the number trajectory in each grid box

res = zeros(splitNumber, splitNumber); % set the count in each grid to zero
rowspan = (toInitialCondition(index(1)) - fromInitialCondition(index(1)))/splitNumber; % define the row_no. from the range of variable 1
% rowspan
colspan = (toInitialCondition(index(2)) - fromInitialCondition(index(2)))/splitNumber; % define the column_no. from the range of variable 2
% colspan
for i = 1:size(allTrajectories, 1)
    for j = 1:size(allTrajectories{i}, 1)
        row = ceil((allTrajectories{i}(j, index(1)) - fromInitialCondition(index(1)))/rowspan) + 1;
%         row
        col = ceil((allTrajectories{i}(j, index(2)) - fromInitialCondition(index(2)))/colspan) + 1;


            if (row > 0 && row < splitNumber) && (col > 0 && col < splitNumber)               
                res(row, col) = res(row, col) + 1;
            end
    end
end
```
