## Supplementary material for "A Monte Carlo method for *in silico* modeling and visualization of Waddington’s epigenetic landscape with intermediate details": MATLAB source code of our Monte Carlo method and case study models: GetTrajectory.m

```
% 
% The below CLE simulation was modified based on the code from the paper
% Higham DJ. Modeling and simulating chemical reactions. SIAM Rev. 2008;50(2):347-68.
%
% Simple implementation of Euler--Maruyama to simulate the 
% Chemical Langevin Equation for Gardner et al. (2000) model of synthetic toggle switch.
%
%
% CLE_PLOT.M Downloadable from 
%    http://www.maths.strath.ac.uk/~aas96106/algfiles.html
% This is an extended version of cle_mm.m that 
% produces graphical output. 
%


function res = GetTrajectory(initialConditions, vol)

% randn('state',100)

% stoichiometric matrix
% define the stoichiometric matrix V for 4 reactions
V = [1 -1 0 0; 0 0 1 -1];

%%%%%%%%%% Parameters and Initial Conditions %%%%%%%%%
nA = 6.023e23;             % Avagadro's number
% vol = 1e-20;               % volume of system original Higham paper is vol=1e-15
% smaller vol produces larger noise
Y = zeros(2,1);

% set the initial conditions (in number of molecules)
Y(:,1) = initialConditions(1,:)';

% rate constants formulation from k to c
c(1) = 3*(nA*vol); 
c(2) = 1*nA*vol; 
c(3) = 1;
c(4) = 3*(nA*vol);
c(5) = 1;

% simulation time
tfinal = 30;

% stepsize
L = 250;
tau = tfinal/L;    

Yvals = zeros(2,L+1);
Yvals(1,1) = Y(1); % this mean Y(1,1)
Yvals(2,1) = Y(2); % this mean Y(2,1) and so on.

for k = 1:L
%     
     a(1) = c(1)*c(2)^3/(c(2)^3+Y(2)^3);
     
     a(2) = c(3)*Y(1);
     
     a(3) = c(4)*c(2)^3/(c(2)^3+Y(1)^3);
    
     a(4) = c(5)*Y(2);

     d(1) = tau*a(1)  + sqrt(abs(tau*a(1)))*randn;
     d(2) = tau*a(2)  + sqrt(abs(tau*a(2)))*randn;
     d(3) = tau*a(3)  + sqrt(abs(tau*a(3)))*randn;
     d(4) = tau*a(4)  + sqrt(abs(tau*a(4)))*randn;
     
     % Y(t+tau)= Y(t) + tau*(aj*vj) + sqrt(tau*aj)*Zj*vj
     %                  for j=1 to M=4
     Y = Y + d(1)*V(:,1) + d(2)*V(:,2) + d(3)*V(:,3)+ d(4)*V(:,4);   

     Yvals(:,k+1) = Y;
end

res = Yvals';
```
