## Supplementary material for "A Monte Carlo method for *in silico* modeling and visualization of Waddington’s epigenetic landscape with intermediate details": MATLAB source code of our Monte Carlo method and case study models: Setting_and_running_CLE.m

```
%% Setting_and_running_CLE.m implements the algorithm to calculate the potential landscape, plot the figures in 3D view and Top view, 
% and finally record the time it takes.

%% Author: Xiaomeng Zhang and Ket Hing Chong
% School of Computer Science and Engineering
% Nanyang Technological University
% Singapore
%
% Last revision: 7 Sept 2017


clear all
close all

addpath('../common_code_CLE'); % to access the common MATLAB files
tic

% InitParams;
variableNames = {'u', 'v'};

index = [1 2];

% Avogadro's number
nA = 6.023e23;             
global vol;
vol = 1e-20;  % original vol = 1e-15

range_max = 4;
range_min = 0;
% set the range of the space of interest (you can get maximum range from time course simulations)
initialRange = [zeros(1,2)+range_min;   % set the minimum range
                zeros(1,2)+range_max]; % set the maximum range


trajectoryNumber = 100000;

splitNumber = 100;
            
fromInitialCondition = min(initialRange);
toInitialCondition = max(initialRange);


% Get all random initial conditions
for j = 1:length(fromInitialCondition)
%   get each random initial conditions within the state space (in number of molecules)
    res(:,j)=nA*vol*(rand(trajectoryNumber, 1)*(toInitialCondition(j)-fromInitialCondition(j))+fromInitialCondition(j));
end

% set the initial conditions as obtained from above
initialConditions = res;

% Call the DrawLandscape function to draw landscape:
DrawLandscape;
```
