## Supplementary material for "A Monte Carlo method for *in silico* modeling and visualization of Waddington’s epigenetic landscape with intermediate details": MATLAB source code of our Monte Carlo method and case study models: GetTrajectory.m

```
% 
% The below CLE simulation was modified based on the code from the paper
% Higham DJ. Modeling and simulating chemical reactions. SIAM Rev. 2008;50(2):347-68.
%
% Simple implementation of Euler--Maruyama to simulate the 
% Chemical Langevin Equation for the Li and Wang (2015) model of cancer attractor.
%
%
% CLE_PLOT.M Downloadable from 
%    http://www.maths.strath.ac.uk/~aas96106/algfiles.html
% This is an extended version of cle_mm.m that 
% produces graphical output. 
%

function res = GetTrajectory(initialCondition, vol)

% updated on: 23 August 2016

% randn('state',100)

% define the stoichiometric matrix V for 22 reactions
% each column of V represents one reaction
%    v1 v2  v3 v4 v5 v6 v7 v8 v9 v10 v11 v12 v13 v14 v15 v16 v17 v18 v19
%    v20 v21 v22 
% each row represents one molecular species     
V = [1  1   -1  0   0   0   0   0   0   0   0   0   0   0   0   0   0   0   0   0   0   0;
    0   0   0   1   1   -1  0   0   0   0   0   0   0   0   0   0   0   0   0   0   0   0;
    0   0   0   0   0   0   1   1   1   -1  0   0   0   0   0   0   0   0   0   0   0   0;
    0   0   0   0   0   0   0   0   0   0   1   1   1   -1  0   0   0   0   0   0   0   0;
    0   0   0   0   0   0   0   0   0   0   0   0   0   0   1   1   1   -1  0   0   0   0;
    0   0   0   0   0   0   0   0   0   0   0   0   0   0   0   0   0   0   1   1   1   -1];

%%%%%%%%%% Parameters and Initial Conditions %%%%%%%%%
nA = 6.023e23;             % Avagadro's number
% vol = 1e-20               % volume of system original Higham paper is vol=1e-15
Y = zeros(6,1);


Y(:,1) = initialCondition(1,:)';

% rate constants
a=0.5*nA*vol;b=0.1*nA*vol;S=0.5*nA*vol;n=4;k=1;
sa=0.8*nA*vol;

c(1) = sa; c(2) = b;  c(3)= k; 

c(4)=a;c(5)=b;c(6)=k;

c(7)=sa;c(8)=b;c(9)=b;c(10)=k;

c(11)=a;c(12)=b;c(13)=b;c(14)=k;

c(15)=sa;c(16)=b;c(17)=b;c(18)=k;

c(19)=a;c(20)=a;c(21)=b;c(22)=k;


tfinal = 30;
L = 250;
tau = tfinal/L;    % stepsize

Yvals = zeros(6,L+1);
Yvals(1,1) = Y(1); % this mean Y(1,1)
Yvals(2,1) = Y(2); % this mean Y(2,1) and so on.
Yvals(3,1) = Y(3);
Yvals(4,1) = Y(4);
Yvals(5,1) = Y(5);
Yvals(6,1) = Y(6);

for k = 1:L
     a(1) = c(1)*Y(1)^n/(S^n+Y(1)^n);
     
     a(2) = c(2)*S^n/(S^n+Y(2)^n);
    
     a(3) = c(3)*Y(1);
     
     a(4) = c(4)*Y(1)^n/(S^n+Y(1)^n);
     
     a(5) = c(5)*S^n/(S^n+Y(4)^n);
     
     a(6) = c(6)*Y(2);
     
     a(7) = c(7)*Y(3)^n/(S^n+Y(3)^n);
     
     a(8) = c(8)*S^n/(S^n+Y(1)^n);
     
     a(9) = c(9)*S^n/(S^n+Y(4)^n);
     
     a(10) = c(10)*Y(3);
     
     a(11) = c(11)*Y(1)^n/(S^n+Y(1)^n);
     
     a(12) = c(12)*S^n/(S^n+Y(3)^n);
     
     a(13) = c(13)*S^n/(S^n+Y(5)^n);

     a(14) = c(14)*Y(4);
     
     a(15) = c(15)*Y(5)^n/(S^n+Y(5)^n);
     
     a(16) = c(16)*S^n/(S^n+Y(4)^n);
     
     a(17) = c(17)*S^n/(S^n+Y(6)^n);
     
     a(18) = c(18)*Y(5);
     
     a(19) = c(19)*Y(1)^n/(S^n+Y(1)^n);
     
     a(20) = c(20)*Y(3)^n/(S^n+Y(3)^n);
     
     a(21) = c(21)*S^n/(S^n+Y(5)^n);
     
     a(22) = c(22)*Y(6);
     
     d(1) = tau*a(1) + sqrt(abs(tau*a(1)))*randn;
     d(2) = tau*a(2) + sqrt(abs(tau*a(2)))*randn;
     d(3) = tau*a(3) + sqrt(abs(tau*a(3)))*randn;
     d(4) = tau*a(4) + sqrt(abs(tau*a(4)))*randn;
     d(5) = tau*a(5) + sqrt(abs(tau*a(5)))*randn;
     d(6) = tau*a(6) + sqrt(abs(tau*a(6)))*randn;
     d(7) = tau*a(7) + sqrt(abs(tau*a(7)))*randn;
     d(8) = tau*a(8) + sqrt(abs(tau*a(8)))*randn;
     d(9) = tau*a(9) + sqrt(abs(tau*a(9)))*randn;
     d(10) = tau*a(10) + sqrt(abs(tau*a(10)))*randn;
     d(11) = tau*a(11) + sqrt(abs(tau*a(11)))*randn;
     d(12) = tau*a(12) + sqrt(abs(tau*a(12)))*randn;
     d(13) = tau*a(13) + sqrt(abs(tau*a(13)))*randn;
     d(14) = tau*a(14) + sqrt(abs(tau*a(14)))*randn;
     d(15) = tau*a(15) + sqrt(abs(tau*a(15)))*randn;
     d(16) = tau*a(16) + sqrt(abs(tau*a(16)))*randn;
     d(17) = tau*a(17) + sqrt(abs(tau*a(17)))*randn;
     d(18) = tau*a(18) + sqrt(abs(tau*a(18)))*randn;
     d(19) = tau*a(19) + sqrt(abs(tau*a(19)))*randn;
     d(20) = tau*a(20) + sqrt(abs(tau*a(20)))*randn;
     d(21) = tau*a(21) + sqrt(abs(tau*a(21)))*randn;
     d(22) = tau*a(22) + sqrt(abs(tau*a(22)))*randn;
     
     % CLE definition as in Eqn (12) in Higham paper
     % Y(t+tau)= Y(t) + tau*(aj*vj) + sqrt(tau*aj)*Zj*vj
     %                  for j=1 to M=22
     Y = Y + d(1)*V(:,1) + d(2)*V(:,2)  + d(3)*V(:,3)+ d(4)*V(:,4)+ d(5)*V(:,5)+ d(6)*V(:,6)+ d(7)*V(:,7)+ d(8)*V(:,8)+ d(9)*V(:,9)+ ...
         + d(10)*V(:,10)+ d(11)*V(:,11)+ d(12)*V(:,12)+ d(13)*V(:,13)+ d(14)*V(:,14)+ d(15)*V(:,15)+ d(16)*V(:,16)+ d(17)*V(:,17)+ d(18)*V(:,18)+ d(19)*V(:,19)+ ...
         + d(20)*V(:,20)+ d(21)*V(:,21)+ d(22)*V(:,22);

     Yvals(:,k+1) = Y;
end

res = Yvals';
```
