## Supplementary material for "A Monte Carlo method for *in silico* modeling and visualization of Waddington’s epigenetic landscape with intermediate details": MATLAB source code of our Monte Carlo method and case study models: README.txt

----------------------------------------------------------------------------------------------------
--- Documentation for drawing Waddington's epigenetic Landscape in stochastic approach using CLE----
----------------------------------------------------------------------------------------------------

Table of Contents
====================
- Introduction
- Usage
- Additional Information


Introduction
============

The algorithm of the Monte Carlo method for quantifying Waddington's epigenetic landscape was implemented in the MATLAB code distributed in different folders for two gene regulatory networks as discussed in our paper.This README explain how you can run the codes in MATLAB to generate the Waddington's epigenetic landscape for stochastic approach using CLE.The CLE implementation was adopted from Higham DJ. titled: 
Modeling and simulating chemical reactions. SIAM Rev. 2008;50(2):347-68.

This MATLAB code package can be modified according to your gene regulatory network in ODEs. The ODEs can be converted to CLE with the different rate constants and stoichiometric matrix V which have to be defined in the Equations.m file.
For details on how to use the code, please refer to Online Supplementary Information. 


Usage
=====
Open the model folder you want to run and plot the Waddington's epigenetic landscape.
Each folder contains 2 MATLAB codes:
1. Setting_and_running_CLE.m 
2. GetTrajectory.m


To run and plot the Waddington's epigenetic landscape, open the first m file (Setting_and_running_CLE.m) in MATLAB editor and click the Run button in the menu bar.

Alternatively, you can run and plot the Waddington's epigenetic landscape in the MATLAB command line.
Type the filename and press Enter at the command line as below:
>>  Setting_and_running_CLE

After the calculation finished with 100,000 time course simulations, you will obtain two graphs: one for 3D view and the other one for Top view of the landscape.


Additional Information
======================

The MATLAB codes were written by Xiaomeng Zhang and Ket Hing Chong. 

This MATLAB code package can be modified according to your gene regulatory network in CLEs, which has to be defined in the GetTrajectory.m file.
Please refer to Online Supplementary Information. 

For any question or comments, please contact Jie Zheng <>
