## Supplementary material for "A Monte Carlo method for *in silico* modeling and visualization of Waddington’s epigenetic landscape with intermediate details": MATLAB source code of our Monte Carlo method and case study models: DrawLandscape.m

```
% Get a randomized initial conditions for the trajectoryNumber defined
initialConditions = GetAllInitialConditions(fromInitialCondition, toInitialCondition, trajectoryNumber);

allTrajectories = GetAllTrajectories(initialConditions, t);

%%

% Plot 3D view of the potential landscape
figure;

% Get the probability distributions
positionProbabilities = GetPositionProbabilities(fromInitialCondition, toInitialCondition, index, splitNumber, allTrajectories);

% Normalized Probability
normalized_Prob=positionProbabilities./(sum(sum(positionProbabilities)));

% U = - ln P
Neg_lnP=-log(normalized_Prob); 

[x y] = meshgrid(linspace(fromInitialCondition(index(1)), toInitialCondition(index(1)), splitNumber), ...
    linspace(fromInitialCondition(index(2)), toInitialCondition(index(2)), splitNumber));

% plot the 3D view of the potential landscape
surf(x,y,Neg_lnP);

% label the axes
xlabel(variableNames(index(2)));
ylabel(variableNames(index(1)));
zlabel('U')

% set the x and y axes
axis([range_min range_max range_min range_max]);

% 3D view
view(-13,66); 
box on;

% set the color bar 
colorbar;

%%

% Plot Top view of the potential landscape
figure;

surf(x,y,Neg_lnP);
xlabel(variableNames(index(2)));
ylabel(variableNames(index(1)));
zlabel('U')

axis([range_min range_max range_min range_max]);

% Top view
view(0,90);
box on;
colorbar;

% show how long it takes to calculate and plot the landscape
disp('Calculation and plotting finished.')
duration=toc/(60*60);
disp(['It took ' num2str(duration) ' hours'])
```
