## Supplementary material for "A Monte Carlo method for *in silico* modeling and visualization of Waddington’s epigenetic landscape with intermediate details": MATLAB source code of our Monte Carlo method and case study models: GetAllInitialConditions.m

```
function res = GetAllInitialConditions(fromInitialCondition, toInitialCondition, trajectoryNumber)

res = zeros(trajectoryNumber, length(fromInitialCondition));
for j = 1:length(fromInitialCondition)
    % Set the initial conditions randomly in the state space
    res(:, j) = rand(trajectoryNumber, 1)*(toInitialCondition(j)-fromInitialCondition(j)) ...
    + fromInitialCondition(j);
end
```
