## Supplementary material for "A Monte Carlo method for *in silico* modeling and visualization of Waddington’s epigenetic landscape with intermediate details": MATLAB source code of our Monte Carlo method and case study models: GetAllTrajectories.m

```
function res = GetAllTrajectories(initialConditions, t)
% Use a loop to calculate all the trajectories or time course simulations
disp('Start calculations ...')
res = cell(size(initialConditions, 1),1);
for i = 1:size(initialConditions, 1)
    % Show the progress of the calculation 
    disp(i);
    % Calculate the time course simulation according to each initial
    % conditions
    res{i} = GetOneTrajectory(t, initialConditions(i, :));
end
```
