## Supplementary material for "A Monte Carlo method for *in silico* modeling and visualization of Waddington’s epigenetic landscape with intermediate details": MATLAB source code of our Monte Carlo method and case study models: equations.m

```
function dy = equations	( t, y, params)
% set the initial size of ODEs
dy=zeros(3,1);

alpha1=0.1;alpha2=3;alpha3=3;
beta1=3;beta2=1;beta3=1;
K1=0.5; K2=0.5;K3=0.5;
n1=8; n2=8; n3=8;

% Define the equations according to the ODEs  
% Model equations given in
% Ferrell James�E, Tsai Tony�Y-C, Yang Q. Modeling the Cell Cycle: Why Do Certain Circuits Oscillate? Cell. 2011;144(6):874-85.
% CDK1a
dy(1)=alpha1-beta1*y(1)*y(3)^n1/(K1^n1+y(3)^n1);

%Plk1a
dy(2)=alpha2*(1-y(2))*y(1)^n2/(K2^n2+y(1)^n2)-beta2*y(2);

%APCa
dy(3)=alpha3*(1-y(3))*y(2)^n3/(K3^n3+y(2)^n3)-beta3*y(3);

end
```
