## Supplementary material for "A Monte Carlo method for *in silico* modeling and visualization of Waddington’s epigenetic landscape with intermediate details": MATLAB source code of our Monte Carlo method and case study models: Setting_and_running.m

```
%% Setting_and_running.m implements the algorithm to calculate the potential landscape, plot the figures in 3D view and Top view, 
% and finally record the time it takes.

% Author: Xiaomeng Zhang and Ket Hing Chong
% School of Computer Science and Engineering
% Nanyang Technological University
% Singapore
%
% Last revision: 7 Sept 2017

clear all;
close all;

addpath('../common_code'); % to access the common MATLAB files
% time in clock to record the start time
tic;

% define the variables name using the order as in ODEs equations
% For the example 5, there are 3 variables
variableNames = {'CDK1a', 'Plk1a','APCa'};


range_max=2;
range_min=0;
% set the range of the space of interest (you can get maximum range from time course simulations)
initialRange = [zeros(1,3)+range_min;    % set the minimum range
               zeros(1,3)+range_max]; % set the maximum range
           
index = [1 2]; % the index of the two variables of interest to view the phase plane (1 for CDK1a and 2 for Plk1a)

% t define the numerical integration interval
t = 0:0.1:30; % set the max end time

fromInitialCondition = min(initialRange);
toInitialCondition = max(initialRange);

% set the number of trajectories for time course simulations 
trajectoryNumber = 100000; 

% set the number of grid box size (100 x 100)
splitNumber = 100;

% Call the DrawLandscape function to draw landscape:
DrawLandscape;
```
