## Supplementary material for "A Monte Carlo method for *in silico* modeling and visualization of Waddington’s epigenetic landscape with intermediate details": MATLAB source code of our Monte Carlo method and case study models: equations.m

```
function dy = equations( t, y)
% set the initial dimension of the model
dy=zeros(2,1);

% define the model equations according to ODEs
% We used the model equations given in
% Chong KH, Samarasinghe S, Kulasiri D, Zheng J. Computational techniques in mathematical modelling of biological switches. 
% In: Weber T, McPhee MJ, Anderssen RS, editors. 21st International Congress on Modelling and Simulation (MODSIM 2015); Gold Coast, Australia 2015. p. 578-84.
dy(1)=3/(1+y(2)^3)-y(1);
dy(2)=3/(1+y(1)^3)-y(2);
end
```
