## Supplementary material for "A Monte Carlo method for *in silico* modeling and visualization of Waddington’s epigenetic landscape with intermediate details": MATLAB source code of our Monte Carlo method and case study models: equations.m

```
function dy = equations	( t, y)
% 52-dimensional dynamical system with two attractors
%  model equations taken from Li, C. and Wang, J. (2013) 
%  Quantifying cell fate decisions for differentiation and reprogramming of a human stem cell network: landscape and biological paths.
%  PLoS Comp. Biol.;9(8):e1003165.

% set the initial size of ODEs
dy=zeros(52,1);

% Define the equations according to the ODEs  

% #1 CDX2'
dy(1)=-y(1)+0.37*y(1)^3/(0.5^3+y(1)^3)+0.5*0.5^3/(0.5^3+y(45)^3)+0.5*0.5^3/(0.5^3+y(44)^3);
% #2 PBX1'
dy(2)=-y(2)+0.37*y(44)^3/(0.5^3+y(44)^3);
% #3 GATA6'
dy(3)=-y(3)+0.37*y(3)^3/(0.5^3+y(3)^3)+0.5*0.5^3/(0.5^3+y(14)^3)+0.5*0.5^3/(0.5^3+y(45)^3)+0.5*0.5^3/(0.5^3+y(15)^3)+0.5*0.5^3/(0.5^3+y(44)^3)+0.5*0.5^3/(0.5^3+y(12)^3);
% #4 E2F4'
dy(4)=-y(4);
% #5 TDGF1'
dy(5)=-y(5)+0.37*y(44)^3/(0.5^3+y(44)^3)+0.37*y(45)^3/(0.5^3+y(45)^3)+0.37*y(47)^3/(0.5^3+y(47)^3);
% #6 FOXA2'
dy(6)=-y(6)+0.37*y(50)^3/(0.5^3+y(50)^3)+0.37*y(6)^3/(0.5^3+y(6)^3)+0.37*y(7)^3/(0.5^3+y(7)^3)+0.5*0.5^3/(0.5^3+y(8)^3);
% #7 SOX17'
dy(7)=-y(7)+0.5*0.5^3/(0.5^3+y(48)^3)+0.5*0.5^3/(0.5^3+y(45)^3);
% #8 OCT4FOXD3'
dy(8)=-y(8)+0.37*y(45)^3/(0.5^3+y(45)^3)+0.37*y(50)^3/(0.5^3+y(50)^3);
% #9 FOXA1'
dy(9)=-y(9)+0.37*y(9)^3/(0.5^3+y(9)^3)+0.37*y(50)^3/(0.5^3+y(50)^3)+0.5*0.5^3/(0.5^3+y(8)^3);
% #10 AFP'
dy(10)=-y(10)+0.37*y(9)^3/(0.5^3+y(9)^3)+0.37*y(6)^3/(0.5^3+y(6)^3);
% #11 SUMO1'
dy(11)=-y(11)+0.37*y(44)^3/(0.5^3+y(44)^3)+0.37*y(4)^3/(0.5^3+y(4)^3);
% #12 GATA4'
dy(12)=-y(12)+0.37*y(11)^3/(0.5^3+y(11)^3)+0.37*y(13)^3/(0.5^3+y(13)^3)+0.5*0.5^3/(0.5^3+y(44)^3)+0.5*0.5^3/(0.5^3+y(45)^3);
% #13 PIAS1'
dy(13)=-y(13)+0.37*y(13)^3/(0.5^3+y(13)^3)+0.37*y(4)^3/(0.5^3+y(4)^3);
% #14 PRDM14'
dy(14)=-y(14)+0.37*y(44)^3/(0.5^3+y(44)^3)+0.37*y(45)^3/(0.5^3+y(45)^3)+0.37*y(47)^3/(0.5^3+y(47)^3);
% #15 LMCD1'
dy(15)=-y(15)+0.37*y(44)^3/(0.5^3+y(44)^3)+0.5*0.5^3/(0.5^3+y(22)^3);
% #16 T'
dy(16)=-y(16)+0.5*0.5^3/(0.5^3+y(45)^3)+0.5*0.5^3/(0.5^3+y(44)^3)+0.5*0.5^3/(0.5^3+y(14)^3)+0.5*0.5^3/(0.5^3+y(4)^3);
% #17 CSH1'
dy(17)=-y(17)+0.5*0.5^3/(0.5^3+y(14)^3);
% #18 GATA2'
dy(18)=-y(18)+0.5*0.5^3/(0.5^3+y(47)^3)+0.5*0.5^3/(0.5^3+y(44)^3)+0.5*0.5^3/(0.5^3+y(19)^3)+0.5*0.5^3/(0.5^3+y(11)^3);
% #19 PIASy'
dy(19)=-y(19)+0.37*y(13)^3/(0.5^3+y(13)^3);
% #20 GATA3'
dy(20)=-y(20)+0.5*0.5^3/(0.5^3+y(44)^3);
% #21 hCGa'
dy(21)=-y(21)+0.37*y(18)^3/(0.5^3+y(18)^3)+0.37*y(20)^3/(0.5^3+y(20)^3)+0.37*y(22)^3/(0.5^3+y(22)^3)+0.5*0.5^3/(0.5^3+y(45)^3);
% #22 cAMP'
dy(22)=-y(22);
% #23 SP1'
dy(23)=-y(23)+0.37*y(4)^3/(0.5^3+y(4)^3)+0.37*y(23)^3/(0.5^3+y(23)^3)+0.37*y(41)^3/(0.5^3+y(41)^3)+0.37*y(30)^3/(0.5^3+y(30)^3)+0.5*0.5^3/(0.5^3+y(40)^3)+0.5*0.5^3/(0.5^3+y(25)^3);
% #24 hCGb'
dy(24)=-y(24)+0.37*y(23)^3/(0.5^3+y(23)^3)+0.37*y(25)^3/(0.5^3+y(25)^3)+0.37*y(22)^3/(0.5^3+y(22)^3)+0.5*0.5^3/(0.5^3+y(45)^3);
% #25 SP3'
dy(25)=-y(25)+0.5*0.5^3/(0.5^3+y(13)^3)+0.5*0.5^3/(0.5^3+y(11)^3);
% #26 BMP2K'
dy(26)=-y(26)+0.37*y(28)^3/(0.5^3+y(28)^3);
% #27 BMP2'
dy(27)=(-1)*y(27)+0.37*y(26)^3/(0.5^3+y(26)^3)+0.5*0.5^3/(0.5^3+y(47)^3)+0.5*0.5^3/(0.5^3+y(44)^3);
% #28 HNF4A'
dy(28)=-y(28)+0.37*y(3)^3/(0.5^3+y(3)^3)+0.37*y(29)^3/(0.5^3+y(29)^3);
% #29 bCAT'
dy(29)=-y(29);
%#30 CEBP'
dy(30)=-y(30)+0.5*0.5^3/(0.5^3+y(4)^3)+0.5*0.5^3/(0.5^3+y(23)^3);
%#31 FOXO1A'
dy(31)=-y(31)+0.37*y(44)^3/(0.5^3+y(44)^3)+0.37*y(45)^3/(0.5^3+y(45)^3)+0.37*y(47)^3/(0.5^3+y(47)^3);
%#32 LEF1'
dy(32)=-y(32)+0.5*0.5^3/(0.5^3+y(45)^3)+0.5*0.5^3/(0.5^3+y(44)^3);
%#33 LEF1bCat'
dy(33)=(-1)*y(33)+0.37*y(32)^3/(0.5^3+y(32)^3)+0.37*y(29)^3/(0.5^3+y(29)^3);
%#34 Mad'
dy(34)=-y(34);
%#35 MadMax'
dy(35)=-y(35)+0.37*y(34)^3/(0.5^3+y(34)^3)+0.37*y(36)^3/(0.5^3+y(36)^3);
%#36 MAX'
dy(36)=-y(36);
%#37 MYC'
dy(37)=-y(37)+0.37*y(33)^3/(0.5^3+y(33)^3)+0.37*y(14)^3/(0.5^3+y(14)^3)+0.37*y(4)^3/(0.5^3+y(4)^3)+0.5*0.5^3/(0.5^3+y(38)^3);
%#38 TCF3'
dy(38)=(-1)*y(38)+0.37*y(4)^3/(0.5^3+y(4)^3);
%#39 MycMax'
dy(39)=(-1)*y(39)+0.37*y(36)^3/(0.5^3+y(36)^3)+0.37*y(37)^3/(0.5^3+y(37)^3);
%#40 MYCSP1'
dy(40)=(-1)*y(40)+0.37*y(23)^3/(0.5^3+y(23)^3)+0.37*y(37)^3/(0.5^3+y(37)^3);
%#41 NFYA'
dy(41)=-y(41)+0.37*y(4)^3/(0.5^3+y(4)^3)+0.37*y(47)^3/(0.5^3+y(47)^3);
%#42 SALL4'
dy(42)=-y(42)+0.37*y(32)^3/(0.5^3+y(32)^3)+0.37*y(44)^3/(0.5^3+y(44)^3)+0.37*y(47)^3/(0.5^3+y(47)^3);
%#43 ZNF206'
dy(43)=-y(43)+0.37*y(44)^3/(0.5^3+y(44)^3);
%#44 NANOG'
dy(44)=-y(44)+0.37*y(44)^3/(0.5^3+y(44)^3)+0.37*y(46)^3/(0.5^3+y(46)^3)+0.37*y(2)^3/(0.5^3+y(2)^3)+0.37*y(48)^3/(0.5^3+y(48)^3)+0.37*y(49)^3/(0.5^3+y(49)^3)+0.5*0.5^3/(0.5^3+y(3)^3);
%#45 OCT4'
dy(45)=-y(45)+0.37*y(44)^3/(0.5^3+y(44)^3)+0.37*y(46)^3/(0.5^3+y(46)^3)+0.5*0.5^3/(0.5^3+y(1)^3);
%#46 OCT4Sox2'
dy(46)=-y(46)+0.37*y(45)^3/(0.5^3+y(45)^3)+0.37*y(47)^3/(0.5^3+y(47)^3);
%#47 SOX2'
dy(47)=-y(47)+0.37*y(44)^3/(0.5^3+y(44)^3)+0.37*y(46)^3/(0.5^3+y(46)^3);
%#48 ZIC3'
dy(48)=-y(48)+0.37*y(44)^3/(0.5^3+y(44)^3)+0.37*y(47)^3/(0.5^3+y(47)^3);
%#49 KLF4'
dy(49)=-y(49)+0.37*y(44)^3/(0.5^3+y(44)^3)+0.37*y(45)^3/(0.5^3+y(45)^3)+0.37*y(47)^3/(0.5^3+y(47)^3);
%#50 FOXD3'
dy(50)=-y(50)+0.37*y(44)^3/(0.5^3+y(44)^3)+0.37*y(45)^3/(0.5^3+y(45)^3)+0.37*y(47)^3/(0.5^3+y(47)^3)+0.37*y(4)^3/(0.5^3+y(4)^3);
%#51 ZFP42'
dy(51)=-y(51)+0.37*y(44)^3/(0.5^3+y(44)^3)+0.37*y(45)^3/(0.5^3+y(45)^3);
%#52 GDF3'
dy(52)=-y(52)+0.37*y(47)^3/(0.5^3+y(47)^3);

end
```
