## Supplementary material for "A Monte Carlo method for *in silico* modeling and visualization of Waddington’s epigenetic landscape with intermediate details": MATLAB source code of our Monte Carlo method and case study models: Setting_and_running.m

```
%% Setting_and_running.m implements the algorithm to calculate the potential landscape, plot the figures in 3D view and Top view, 
% and finally record the time it takes.

% Author: Xiaomeng Zhang and Ket Hing Chong
% School of Computer Science and Engineering
% Nanyang Technological University
% Singapore
%
% Last revision: 7 Sept 2017

clear all;
close all;

addpath('../common_code'); % to access the common MATLAB files

% time in clock to record the start time
tic;

% define the variables name using the order as in ODEs equations
% For the example 5, there are 52 variables
variableNames = {'CDX2', 'PBX1', 'GATA6', 'E2F4','TDGF1','FOXA2','SOX17','OCT4FOXD3','FOXA1','AFP',...
              'SUMO1','GATA4','PIAS1','PRDM14', 'LMCD1', 'T', 'CSH1', 'GATA2','PIASy', 'GATA3',...
              'hCGa','CAMP','SP1','hCGb','SP3','BMP2K','BMP2','HNF4A','bCAT','CEBP', ...
              'FOXO1A','LEF1','LEF1bCat','Mad','MadMax','MAX','MYC','TCF3','MycMax','MYCSP1', ...
              'NFYA','SALL4','ZNF206','NANOG','OCT4','OCT4Sox2','SOX2','ZIC3','KLF4','FOXD3', ...
              'ZFP42','GDF3'};


range_max=3.5;
range_min=0;
% set the range of the space of interest (you can get maximum range from time course simulations)
initialRange = [zeros(1,52)+range_min;    % set the minimum range
               zeros(1,52)+range_max]; % set the maximum range

index = [44 3]; % the index of the two variables of interest to view the phase plane (44 for Nanog and 3 for GATA6)

% t define the numerical integration interval
t = 0:0.1:30; % set the max end time

fromInitialCondition = min(initialRange);
toInitialCondition = max(initialRange);

% set the number of trajectories for time course simulations 
trajectoryNumber = 100000; 

% set the number of grid box size (100 x 100)
splitNumber = 100;


% Call the DrawLandscape function to draw landscape:
DrawLandscape;
```
