## Supplementary material for "A Monte Carlo method for *in silico* modeling and visualization of Waddington’s epigenetic landscape with intermediate details": MATLAB source code of our Monte Carlo method and case study models: equations.m

```
function dy = equations	( t, y, params)

% set the initial size of ODEs
dy=zeros(6,1);

a=0.5;
b=0.1; 
k=1;
S=0.5;
n=4;
sa1=0.8;
sa2=0.8;

% Define the equations according to the ODEs  
% We used model equations given in
% Li C, Wang J. Quantifying the Landscape for Development and Cancer from a Core Cancer Stem Cell Circuit. Cancer Res. 2015:canres. 0079.2015.
% P53
dy(1)=sa1*y(1)^n/(S^n+y(1)^n)+b*S^n/(S^n+y(2)^n)-k*y(1);

%Mdm2
dy(2)=a*y(1)^n/(S^n+y(1)^n)+b*S^n/(S^n+y(4)^n)-k*y(2);

%Oct4
dy(3)=sa2*y(3)^n/(S^n+y(3)^n)+b*S^n/(S^n+y(1)^n)+b*S^n/(S^n+y(4)^n)-k*y(3);

%miR145
dy(4)=a*y(1)^n/(S^n+y(1)^n)+b*S^n/(S^n+y(3)^n)+b*S^n/(S^n+y(5)^n)-k*y(4);

%Zeb
dy(5)=sa2*y(5)^n/(S^n+y(5)^n)+b*S^n/(S^n+y(4)^n)+b*S^n/(S^n+y(6)^n)-k*y(5);

%miR200
dy(6)=a*y(1)^n/(S^n+y(1)^n)+a*y(3)^n/(S^n+y(3)^n)+b*S^n/(S^n+y(5)^n)-k*y(6);
end
```
