## Supplementary material for "A Monte Carlo method for *in silico* modeling and visualization of Waddington’s epigenetic landscape with intermediate details": MATLAB source code of our Monte Carlo method and case study models: equations.m

```
function dy = equations	( t, y, params)
% Example 2 Two-dimensional dynamical system with four attractors
% Zhou JX, Aliyu M, Aurell E, Huang S. Quasi-potential landscape in complex multi-stable systems. 
% Journal of The Royal Society Interface. 2012;9(77):3539-53.
% Eqn (5.7)

% set the initial size of ODEs
dy=zeros(2,1);

% Define the equations according to the ODEs  
% x
dy(1)=-1+9*y(1)-2*y(1)^3+9*y(2)-2*y(2)^3; 
% y
dy(2)=1-11*y(1)+2*y(1)^3+11*y(2)-2*y(2)^3; 
end
```
